# ANTIDOTE: A Metadata-Driven Neural Network for Improving CryoEM 3-D Particle Sorting

**DOI:** 10.1101/2025.10.06.680814

**Authors:** Raymond F. Berkeley, Brian D. Cook, Daniel Ji, Armin Foroughi, Yifei He, Maxwell J. Bachochin, Mark A. Herzik

**Author notes:** These authors contributed equally (Raymond F. Berkeley, Brian D. Cook).

## Abstract

Despite the maturation of cryogenic electron microscopy (cryoEM) methodologies, generating high-resolution three-dimensional (3-D) reconstructions from micrographs is a time-intensive process involving iterative rounds of subjective data curation and hyperparameter optimization. Current approaches to particle classification are often unable to remove all low-quality particles from particle stacks, largely due to the low signal-to-noise ratio, the high dimensionality of particle images, and the multiple degrees of freedom associated with each particle’s unknown rotation, orientation, and class assignment. The retention of low-quality particles negatively affects the overall quality of the final EM density and continued efforts to eliminate their deleterious contributions are warranted. Here, we present ANTIDOTE (A Neural network Trained In Deleterious Object deTection and Elimination), a neural network framework that discriminates between constructive and deleterious particles using per-particle metadata generated during 3-D classification in RELION. Using benchmark and real-world cryoEM datasets, we demonstrate that ANTIDOTE paired with RELION 3-D classification achieves higher particle classification accuracy than conventional data processing approaches alone, yielding improvements in reconstruction quality, global and local resolution, and map interpretability while reducing time-consuming hyperparameter optimization. We additionally detail practical use-case scenarios for ANTIDOTE and demonstrate its versatility in increasing particle curation accuracy for high-quality cryoEM reconstruction.

## Introduction

Single-particle cryogenic electron microscopy (cryoEM) has become a leading method for studying the three-dimensional (3-D) structures of biomacromolecules at high resolution.^1,2^ In cryoEM experiments, two-dimensional (2-D) projection images of individual biomacromolecules embedded in a thin layer of vitreous ice are captured in micrographs.^3^ These individual images, referred to as ‘particles’, ideally represent projections of the biomacromolecule being studied at unknown orientations that must be processed in a series of alignment, classification, and signal averaging steps to yield a high-resolution 3-D reconstruction.^4^ The curation of high-quality particle images is challenging due to the inverse nature of the reconstruction problem — reconstructing 3-D objects from 2-D images of unknown orientations — which in turn is confounded by the inherently low signal-to-noise of individual particle images and intrinsic heterogeneity of the particles themselves. The simultaneous removal of deleterious particles (i.e., low-quality particles that do not productively contribute to the final reconstruction), retention of constructive particles (i.e., high-quality particles that represent the biomolecule of interest and contribute productively to the final reconstruction), and accurate assignment of viewing angles is a difficult and incomplete problem that must be solved for every cryoEM dataset.^5^ Because the ground truth in the data and, consequently, the optimal set of data processing parameters for the dataset are not known *a priori*, cryoEM data processing remains a subjective and time-consuming process, relying heavily on the application of expert-guided heuristics and intuition derived from experience.^4,6,7^ The lack of foolproof metrics and the high dimensionality of the data frequently lead to suboptimal parameter selection, typically remedied through extensive hyperparameter and data processing workflow scouting — time and computationally intensive efforts that should be expedited and made more robust.

The standard approach to cryoEM particle curation involves the iterative removal of bad particles using sequential 2-D and 3-D classification steps.^6,8,9^ For example, a practitioner may first elect to sort particles using 2-D classification, where individual particle images are sorted into 2-D class averages that are determined based on signal-averaging particles of similar viewing angles and/or quality. Ideally, particles that are deemed low-quality, or do not represent the biomacromolecule of interest, cluster into 2-D class averages that can be easily removed from the optimal particle stack. The practitioner may then attempt to further remove deleterious particles using 3-D sorting methods (e.g., 3-D classification), which simultaneously generate solutions to each particle’s viewing angle, or Euler angle, with respect to a reference 3-D volume (i.e., class), by similarly sorting particles into 3-D classes in which the Euler angle and class assignment of each particle are allowed to vary until an optimal solution is reached.^4,6–9^ The goal of this process for the purposes of particle curation is to remove all examples of particles that do not constructively represent the biomacromolecule of interest to generate a clean, high-quality particle stack that best embodies a global solution for the system of particles. However, the lack of ground truth and robust validation metrics do not guarantee this solution is a faithful, nor optimal, representation of the biomacromolecule represented in the particle stack.^10–12^

The two most common cryoEM data processing frameworks, RELION and cryoSPARC, both use an expectation-maximization-based framework for particle classification in 2-D and 3-D.^8,9^ This approach is designed to arrive at the best possible solution for all particles without knowing the identity or viewing angle for any given particle at the outset of the particle curation and classification effort. By alternating between classifying subsets of particles to a nascent class average or volume, and measuring their fit to this reference, confidence in the accuracy improves over successive rounds of classification. Simultaneously, with increasing resolution of this class average, or volume, is improved particle classification accuracy and alignment determination. RELION, and later cryoSPARC, extend the expectation-maximization approach with a Bayesian formulation that further improves particle classification performance by marginalizing over all possible solutions during the maximization step in order to account for residual uncertainty and hedge against potential overfitting as particle classification proceeds.^13^

While the conventional approach described above works exceptionally well given the challenges inherent in the particle classification problem, the high dimensionality and low signal-to-noise of the data mean that the decision boundaries made during 2-D and 3-D class sorting frequently overlap.^14^ This frequently leads to the inclusion of deleterious particles in classes that fail to be rejected. These deleterious particles effectively “poison” the dataset, resulting in deconstructive particles in classes that appear to be of high quality. Similar behaviors lead to the loss of constructive particles into classes that are rejected during the classification process, thereby losing the constructive signal that these high-quality particles contain. Consequently, the quality of the final 3-D reconstruction is compromised, leading to lower resolution and lower quality cryoEM maps in the best case and incomplete or misleading structural data in the worst case.^15,16^

In addition to inaccurate particle classification, each particle classification approach involves choosing a set of hyperparameters (e.g., number of classes, uncertainty factor, etc.) whose optimal values are difficult or impossible to determine beforehand.^8,14,17–19^ Although initial values can be predicted, the determination of ideal hyperparameters can represent a significant time investment as incorrect values can further limit the classification accuracy of particles and thereby decrease the quality of the final particle stack. Both limitations of the cryoEM data processing workflow — lack of ground truth knowledge and costly hyperparameter exploration — underscore the need for new approaches that can overcome these challenges to ensure a more accurate and efficient classification process, ultimately improving the reliability and resolution of cryoEM reconstructions.

To address these challenges, we developed a feed-forward artificial neural network-based particle classifier called ANTIDOTE (A Neural network Trained In Deleterious Object deTection and Elimination) that identifies particles that do not constructively contribute to high-quality cryoEM 3-D reconstructions for their elimination. To generate training data for ANTIDOTE, we assembled ground-truth-labeled cryoEM particle datasets of biomacromolecular complexes representing various sizes and symmetries that had been heavily curated using traditional classification approaches. These curated particle stacks were then selectively poisoned with ground-truth deleterious particles that do not represent the biomacromolecule in the associated ground-truth-labeled data. We subsequently used these labeled data to evaluate the performance of particle curation with and without ANTIDOTE and demonstrate ANTIDOTE’s ability to consistently identify high-quality particles using several standard and non-standard cryoEM specimens, including real-world data from the Electron Microscopy Data Bank (EMDB). Importantly, unlike existing methods, ANTIDOTE operates exclusively on particle metadata obtained during 3-D classification in RELION and is agnostic to the identity of the particle itself — it never directly sees the particle image during evaluation — greatly expanding its generalizability to any dataset that can be subjected to RELION 3-D classification. Ultimately, ANTIDOTE enables the rapid generation of a clean particle stack that can be used for generating 3-D reconstructions that exhibit better overall quality metrics than traditional strategies alone.

## Results

### Generation of ground-truth-labeled particle sets for the evaluation of 2-D and 3-D classification efficacy

We initially sought to evaluate the performance of conventional 2-D and 3-D particle classification methodologies to assess limitations in current workflows by generating a standardized set of ground-truth-labeled cryoEM datasets in which constructive and deconstructive particles are labeled (**Fig. 1 a-c, Supplementary Fig. S1-S3**). As a basis for this evaluation, we first prepared six distinct reference cryoEM particle stacks representing different protein complexes covering a range of sizes (e.g., *∼*64 to *∼*700 kDa) and symmetries (e.g., C_1_, C_2_, D_2_, D_7_, and octahedral). Additionally, these sam-ples — rabbit muscle aldolase, mouse heavy chain apoferritin, human hemoglobin, human mortalin-GrpEL1, *Azotobacter vinelandii* nitrogenase molybdenum-iron protein (MoFeP), and *Thermoplasma acidophilum* 20S (T20S) proteasome core — were chosen to represent samples commonly encountered in cryoEM data processing workflows (**Table 1** and **Fig. 1a, c**).^20–24^ Importantly, these labeled datasets were also acquired using various microscope configurations, including 300 and 200 kV illumination sources, and detector systems, spanning Thermo Fisher Scientific Falcon 4(i) and Gatan K2/3 cameras, representing the instrumentation currently available in state-of-the-art cryoEM facilities.^25^ Each dataset was heavily curated to produce the cleanest possible set of high-quality constructive particles, hereafter sometimes referred to as “True” particles, and were verified to generate high-resolution 3-D reconstructions (**Table 1**).^20–24^

**Table 1:** Composition of Poisoned Protein Datasets for 3-D Classification.

| Protein | Poisoning Approach | Total Particles | 3-D Classifications |
| --- | --- | --- | --- |
| Apoferritin | Calibration grid + noise | 925 881 | 192 |
| Aldolase | Calibration grid + noise | 236 125 | 54 |
| Hemoglobin | Template-picked empty grid | 991 705 | 360 |
| MoFeP | Template-picked empty grid | 726 422 | 144 |
| Mortalin | Template-picked empty grid | 208 085 | 216 |
| T20S Proteasome | cryoSPARC 2-D Classification | 119 360 | 108 |

**Figure 1:**
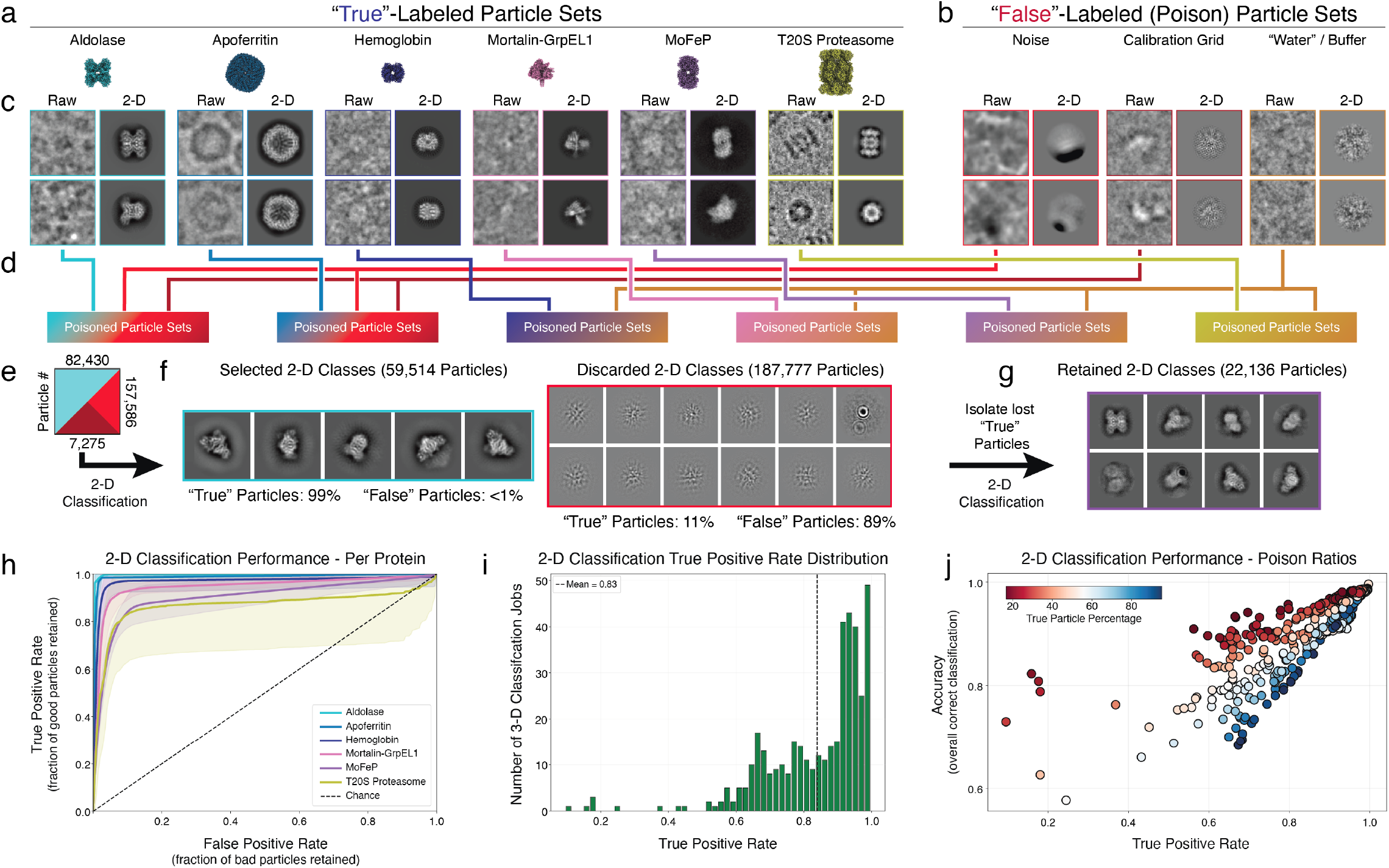
Ground-truth-labeled particle sets indicate expectation-maximization-based 2-D classification is effective but imperfect. **a)** Examples of “True”- and **b)** “False”-labeled (poison) particle sets used for investigating 2-D and 3-D classification efficacy. The sample name and representative image are shown for each test specimen used in this study. **c)** Examples of raw (*left*) and 2-D averaged particles (*right*) for each “True” and “False” specimen are shown. **d)** Combinations of poisoned particle groups (5-500% poison) used in the training and evaluation sets. **e-g**) An example of an aldolase 2-D classification using ground-truth poisoned particles (True=329,722 particles, False=164,861 particles) and representative 2-D class averages, colored by label type, with particle number and particle type percentage shown. **g)** The True particles lost in the discarded 2-D class averages shown in (f) were 2-D classified separately to indicate particle quality. **h)** 2-D classification performance across protein sets. **i)** Histogram showing distribution of TPR values from 2-D classification evaluations. **j)** Accuracy and True Positive Rates (TPR) for cryoSPARC 2-D classifications as a function of the ratio of True (blue) to False (red) particles in the input stack.

We then generated a diverse set of low quality deconstructive particles, hereafter sometimes referred to as “False” or poison particles, sourced from micrographs of a crossed-lines grating replica calibration grid, representing high-contrast contaminating picks, pure noise (i.e., imaged through vacuum) picks, picks resulting from intentional over-picking of micrographs, or template-based picks of “buffer only” micrographs (**Fig. 1b, c**). To ensure that any errors observed in particle curation were due to limitations in 2-D or 3-D sorting and not differences in the images resulting from hardware factors, such as disparities in intensities and contrast transfer function (CTF) values, each True particle stack was exclusively poisoned with False particles collected on the same microscope using the same data collection parameters (see **Methods**) (**Fig. 1d**). This allowed us to generate a panel of poisoned particle stacks that approximates real world mixed experimental particle selections while retaining the ground-truth True and False labels required for an effective classification workflow evaluation (**Fig. 1e-g**). This ground-truth labeling of constructive and deconstructive particles, with True and False labels, respectively, facilitates the use of standard methods for the evaluation of binary classifiers, which we apply for the evaluation of 2-D and 3-D classification performance.^26^

### Assessment of 2-D classification accuracy and limitations using ground-truth-labeled datasets

To evaluate the performance of 2-D classification, we prepared particle stacks containing different ratios of True and False particles for each of the proteins in our dataset (**Table 1; Fig. 1d**). To better reflect the fact that 2-D classification is typically used on uncurated datasets with potentially significant over-picking, we allowed the ratio of True-to-False particles to exceed 1:1 (i.e., *>*100% of the total particle stack were False particles) (**Table 1**). Furthermore, our training sets utilized downsampled particles (e.g., 3-4.5 Å/pixel), consistent with best practices for earlier-stage, less-curated particle stacks generated immediately after particle picking. Each particle stack was subjected to the standard 2-D classification algorithm provided by cryoSPARC, only tuning the inner and outer circular mask diameter to best match the dimensions of the complex in the classification job (**Supplementary Fig. S1-S3**). After each 2-D classification, “good” and “bad” classes were automatically selected using an instance of cryoSPARC’s Reference-Based Auto Select 2D job that had been individually optimized for each biomolecule (**Fig. 1e-f**). After the execution of this standard workflow for 2-D classification and class selection, we treat the Reference-Based Auto Select 2D job as a binary classifier and evaluate the performance of 2-D classification using the ground-truth labels as references.

As a discriminator of constructive and deconstructive particles, cryoSPARC’s 2-D classification appears robust in principle, frequently achieving receiver operating characteristic (ROC) curve area under the curve (AUC) values greater than 0.9 (**Fig. 1h**). To generate ROC curves, we sorted cryoSPARC classes based on the HOG correlations generated for each class in the Reference Based Auto Select 2D job, with classes being perfectly ordered from those most enriched in False particles to those most enriched in True particles. To more accurately reflect the application of 2-D classification in practice, we evaluated the performance of the selected 2-D classes directly using the ground-truth labels. Many of the individual jobs in our analysis achieve accuracies above 80% and the accuracies achieved on a per-protein basis align with structural intuition (**Fig. 1h**). For example, apoferritin, a large, symmetrical, and rigid protein, achieves exceptionally high AUC and accuracy metrics, whereas the mortalin-GrpEL1 complex, which is more dynamic and asymmetrical, is correspondingly more difficult to classify and therefore achieves lower AUC and accuracy metrics. The only protein to achieve a mean AUC of less than 0.9 is the T20S proteasome, which, despite its large size and symmetry, exhibits an unusual inflection at high FPRs in the ROC curve, suggestive of unexpected classification behaviors.

From the ROC curve, we next examined the exact threshold at which cryoSPARC Reference-Based Auto Select 2D discriminates between good and bad particles through 2-D classes. At this cutoff, the True Positive Rate (TPR) across proteins indicates that most particles are generally retained; however, a substantial fraction of jobs still lose a notable number of true particles. On average, *∼*17% of true particles are discarded (*∼*83% retained), and in a non-negligible number of jobs, more than 40% of true particles are lost (**Fig 1i**). Further analysis reveals an interesting trend across the 2-D classification results for the proteins in our panel: as particle stacks become more enriched in false particles, losses in accuracy are largely due to the incorrect rejection of true particles (i.e., constructive particles lost into poor 2-D class averages). This is reflected in the consistently low TPR observed for all 2-D classifications relative to their accuracies, reinforcing our earlier analysis (**Fig. 1j**). For example, a *∼*82K aldolase True particle stack (from EMDB-21023/EMPIAR-10338; see **Methods**) was poisoned with *∼*165K False particles derived from a mixture of calibration grid and noise images (**Fig. 1e**). When subjected to 2-D classification, *∼*59K True particles (72.8% of the True particles in the original particle stack, 99% of the particles in the selected classes) were retained along with only 475 False particles (*<*1% of the False particles in the original particle stack, *<*1% of the particles in the selected classes; **Fig. 1f**). However, 27% of the *∼*187K particles that were in rejected classes were high-quality True-labeled particles (*∼*22K True particles) that could not be recovered through additional 2-D classification steps but could be validated as high-quality constructive particles by manually isolating particles with ground-truth True labels and subjecting the subsequent particle stack to 2-D sorting (**Fig. 1g**).

The observation that more True particles are removed from the particle stack during 2-D class selection as the number of False particles increases has significant implications for particle curation workflows, as it suggests that many constructive particles will be discarded during early 2-D classification runs performed directly after particle picking when deconstructive particles are enriched in particle stacks due to over-picking. This conservative classification behavior, while maintaining high overall accuracy through the effective removal of False particles that dominate the particle stack, leads to an early loss of constructive particles that represents a significant opportunity cost for the reconstruction of high-resolution maps at the end of the data processing pipeline (**Fig. 1f**). The current alternative of carrying a higher False-to-True particle ratio into 3-D sorting and refinement steps to retain more constructive particles presents additional problems that can decrease overall EM map quality.^4,7,15^

### Assessment of 3-D classification accuracy and limitations using ground-truth-labeled datasets

We next sought to use our ground-truth-labeled particle sets to evaluate the performance of particle curation in 3-D using 3-D classification as implemented in RELION. 3-D classification affords the exploration of a wide range of parameters for optimal sorting and can retain lower-populated views that would have been lost during 2-D sorting due to insufficient sampling and poor signal averaging, issues that lead to the misclassification behavior observed in our 2-D evaluations described above (**Fig. 1**).^8^ As with our evaluation of 2-D classification performance, we prepared a similar array of particle stacks with mixed ratios of True and False particles for 3-D classification. To better reflect the fact that 3-D classification is typically used on more curated datasets, we limited the ratio of True-to-False particles to a maximum of 1:1 (i.e., 50-95% of the total particle stack were True particles) (**Table 1**; see **Methods**).^7,8^ Furthermore, similar with our 2-D analyses, each of the particle sets utilizes downsampled particles (e.g., 3-4.5 Å/pixel), consistent with utilization of earlier-stage, less-curated particle stacks. The additional bandwidth afforded by fewer particle ratios was used to diversify the T (Tau fudge factor or regularization parameter), E (E-step or resolution limit), and K (number of classes) 3-D classification parameters. Each particle stack was therefore subjected to multiple RELION 3-D classification jobs using an array of parameters, totaling *>*1000 3-D classification jobs.

Notably, nearly all 3-D classifications in our analysis showed incomplete separation of True and False particles (**Fig. 2a-d, Supplementary Fig. S4**). For example, examination of a 5-class 3-D classification of rabbit muscle aldolase with 32K True particles and *∼*12K False particles derived from a mixture of calibration grid and noise images (see **Methods**) reveals that while many of the classes generate good initial reconstructions of aldolase, False particles are also distributed throughout the classes, making it impossible to separate False particles from True particles solely using class selection (**Fig. 2a**). We then questioned whether the per-particle metadata generated by RELION 3-D classification would possess sufficient discriminative power to distinguish constructive and deconstructive particles, using our ground-truth labels as reference, rather than relying solely on 3-D volume visualization and selection for particle classification.

**Figure 2:**
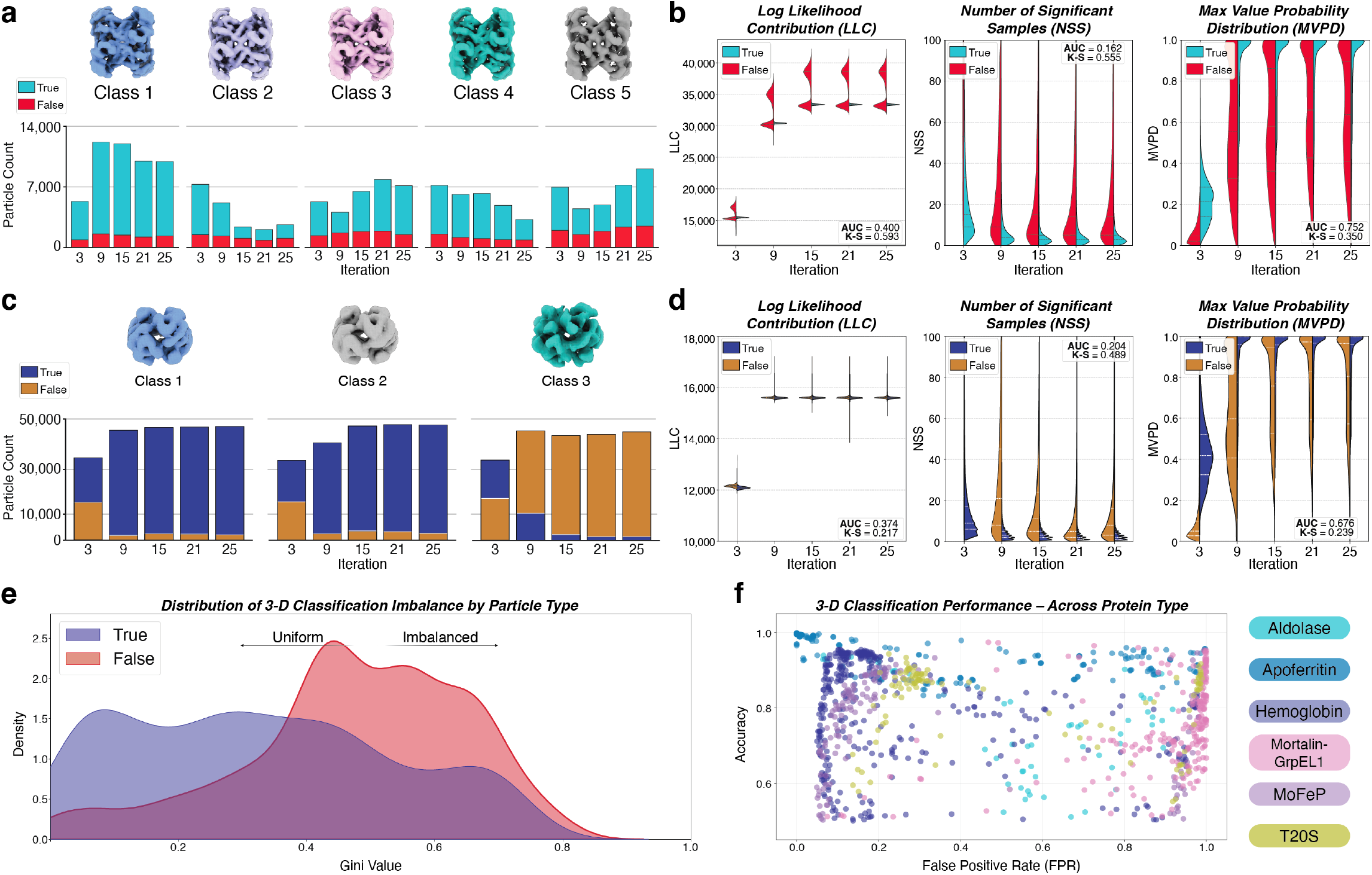
Assessment of expectation-maximization-based 3-D classification efficacy using ground-truth-labeled particle sets. **a)** An example of particle curation using 3-D classification (K=5, T=4, E=6) using a poisoned aldolase dataset (32,000 aldolase particles, 12,125 poison particles) from our ground-truth-labeled evaluation datasets. The True/False ratio for each iteration is shown per class as well as the final 3-D volume. **b)** Violin plots of log-likelihood contribution (LLC), number of significant samples (NSS), and maximum value probability distribution (MVPD) per iteration for True versus False aldolase and poison particles, respectively. Area under the curve (AUC) and Kolmogorov-Smirinov (K-S) statistics are shown as insets. **c)** An example of particle curation using 3-D classification (K=3, T=4) using a poisoned hemoglobin dataset (100,000 hemoglobin particles, 50,000 poison particles) from our ground-truth-labeled evaluation datasets. The True/False ratio for each iteration is shown per class as well as the final 3-D volume. **d)** Violin plots of LLC, NSS, and MVPD per iteration for True versus False hemoglobin and poison particles, respectively. AUC and K-S statistics are shown as insets. **e)** Gini distributions for True- and False-labeled particles across all 3-D classifications assessed using the ground-truth-labeled evaluation datasets. **f)** Accuracy and False Positive Rates (FPR) for RELION 3-D classification for the six proteins in the ground-truth-labeled evaluation datasets.

Each RELION 3-D classification run generates a set of statistics derived from the probability density function describing the fit of each particle to its assigned reference at each iteration of classification: the overall log likelihood (rlnLog-LikeliContribution, LLC), the maximum value of the probability density function (rlnMaxValueProbDistribution, MVPD), and the number of non-noise peaks in the probability density function (rlnNrOfSignificantSamples, NSS). For a constructive particle, the LLC and MVPD values would be expected to be high due to a high-quality fit producing a single large peak in the probability density function. Similarly, the NSS would be expected to be low due to the presence of few additional peaks in the function which would represent alternate fits. For this 3-D classification, which achieves relatively poor accuracy as a particle classifier, the discriminative power of LLC, NSS, and MVPD values at the final iteration of similarly poor, with each metric achieving low AUC and Kolmogorov-Smirinov (K-S) metrics (**Fig. 2b**).

As a point of comparison, we performed the same analysis on a 3-D classification that achieves higher accuracy as a particle classifier. Evaluation of a 3-class classification of hemoglobin (from EMDB-45816/EMPIAR-12743; see **Methods**), reveals three approximately equally populated classes, each with hemoglobin-like features (**Fig. 2c**). Our ground truth labels indicate classes 1 and 2 contain the majority of the True particles (*∼*98% of all True particles, *∼*11% of all False) with class 3 consisting of the remaining *∼*89% of the False particles (2% of the True). Our ground truth labels indicate classes 1 and 2 contain *∼*98% of the True particles (*∼*11% False) with class 3 consisting of *∼*89% of the False particles (2% True). Despite the high accuracy of this 3-D classification run, the LLC, NSS, and MVPD statistics still do not appear to ef-fectively discriminate between constructive and deconstructive particles as standalone classifiers (**Fig. 2d**). Additionally, accuracy in this classification is impacted primarily by the inclusion of False particles in True classes (**Fig. 2c**). In conjunction, our evaluation of the statistics generated by RELION 3-D classification suggests that although the distributions of LLC, NSS, and MVPD trend as expected, insufficient information exists to confidently identify constructive and deconstructive particles in these statistics alone (**Fig. 2b, d, Supplementary Fig. S5**). This conclusion holds for both inaccurate (**Fig. 2a, b**) and accurate (**Fig. 2c, d**) 3-D classifications. This observation implies the existence of additional information in RELION 3-D classification that facilitates the classification of particles in 3-D that may be captured to effect more accurate particle sorting.

### Class Spreading Leads to Misallocation of High-Quality Particles During 3-D Classification

Evaluating the Gini coefficients of True and False particles across all 3-D classifications reveals that True particles are more likely to be broadly distributed across all classes in more even numbers (i.e., lower Gini value) than False particles, which are more imbalanced and frequently distribute into all classes with much greater variability (i.e., higher Gini values) (**Fig. 2e, Supplementary Fig. S6**).^27^ We speculate that this behavior may be due to the presence of distinct features in constructive particles that can be reinforced over iterations of 3-D classification that are not present deconstructive particles, which are more likely to lack consistent averageable features. However, we must note that the Gini coefficients for True particles exhibit a broader distribution than those observed for False particles, likely a result of the quality of the biomacromolecule representing the True particles. Apoferritin, for example, will likely classify into a single True class due to its rigidity and homogeneity, while mortalin-GrpEL1 is more likely to distribute into classes representing distinct conformations (**Supplementary Fig. S6**)

To obtain a more comprehensive picture of the performance of 3-D classification as a particle classifier, we evaluated all 3-D classifications in our dataset by within a binary classification framework to assess particle sorting accuracy. Unlike the evaluation of 2-D classification, where we calculated accuracies using the ground truth labels for particles contained in classes accepted and rejected by cryoSPARC’s Reference-Based Auto Select 2D job, for 3-D classification we instead calculated the accuracy of all possible combinations of class acceptance and rejection using the ground-truth particle labels that retained the highest accuracy value across combinations. To better reflect the application of 3-D classification for particle sorting in the real world, we did not include accuracies achieved if all classes were accepted or rejected (i.e. we enforced the rejection or inclusion of at least one class). Note, since nearly all 3-D classification jobs allocated True particles across all classes, the highest accuracy TPR, but not necessarily accuracy, would result from retaining all classes, regardless of how many False particles contained (**Fig. 2a, c, e**). However, this approach does not represent real-world scenarios since a cryoEM practitioner generally would not select all classes for downstream processing knowing low quality particles existed. Therefore, for each classification job, we evaluated all possible combinations of class retention and removal – selecting the combination of classes that maximizes accuracy while requiring at least one class to be removed. This approach allows us to simulate the optimal decision that would be made by an expert user for each of the 3-D classifications in our evaluation set.

From this analysis, several different trends emerge from our bulk 3-D classification dataset. First, in some cases, such as apoferritin, 3-D classification is robust — it consistently achieves high accuracy and effectively sorts particles into high-quality and low-quality classes across a range of 3-D classification hyperparameters (**Fig. 2f, Supplementary Fig. S4**). Secondly, in numerous cases, 3-D classification struggles to partition particles into classes at all, and instead often bins a large number of True and False particles into a single class. This behavior is often observed with mortalin-GrpEL1 and a subset of the T20S and MoFeP data, and in particle stacks enriched in True particles manifests as a 3-D classification with a high accuracy but also a high FPR due to the automated rejection of sparsely populated classes (**Fig. 2f, Supplementary Fig. S4**). Finally, we also frequently observe the effective classification of False particles into classes that also absorb a significant fraction of True particles. This leads to a low FPR and a variable overall accuracy that depends on the fraction of True particles lost into a low-quality class — a scenario that represents a significant opportunity cost in the final reconstruction. This behavior was often observed with hemoglobin, aldolase, and MoFeP (**Fig. 2f, Supplementary Fig. S4**).

### A feed-forward artificial neural network captures particle behaviors during 3-D classification

Imperfect 2-D and 3-D classification of True and False particles was observed across all specimens in our evaluation, demonstrating a need for additional approaches to exclusively recover True particles. Despite the rich set of statistics generated by RELION for each particle in the 3-D classification run, we noticed that 3-D classification often primarily sorted particles into a small number of classes. We therefore reasoned that there may be a more effective way to leverage 3-D per-particle quality metrics than relying solely on 2-D and 3-D class average selection approaches.

Despite the poor performance of the Bayesian statistical values from 3-D classification as standalone proxies for particle quality (e.g., LLC, NSS, and MVPD), we surmised that information embedded in the classification trajectory of each particle over iterative rounds of 3-D classification, per-particle performance data from 3-D classification encoded in the Bayesian statistics generated by RELION (LLC, NSS, and MVPD), and additional context reflecting the quality of each particle’s class may contain sufficient information to effectively classify particles. We therefore used our ground-truth-labeled particle sets and 3-D classification library to build a framework for training feed-forward artificial neural networks on 3-D classification data and used the large 3-D classification dataset generated by our extensive analysis of poisoned particle stacks to train models for particle classification (**Fig. 3a**). Importantly, the presence of ground-truth False labels in our evaluation data facilitated the generation and evaluation of models less likely to contain biases that may be introduced by any standard approach to particle curation (**Fig. 2a-c**).

**Figure 3:**
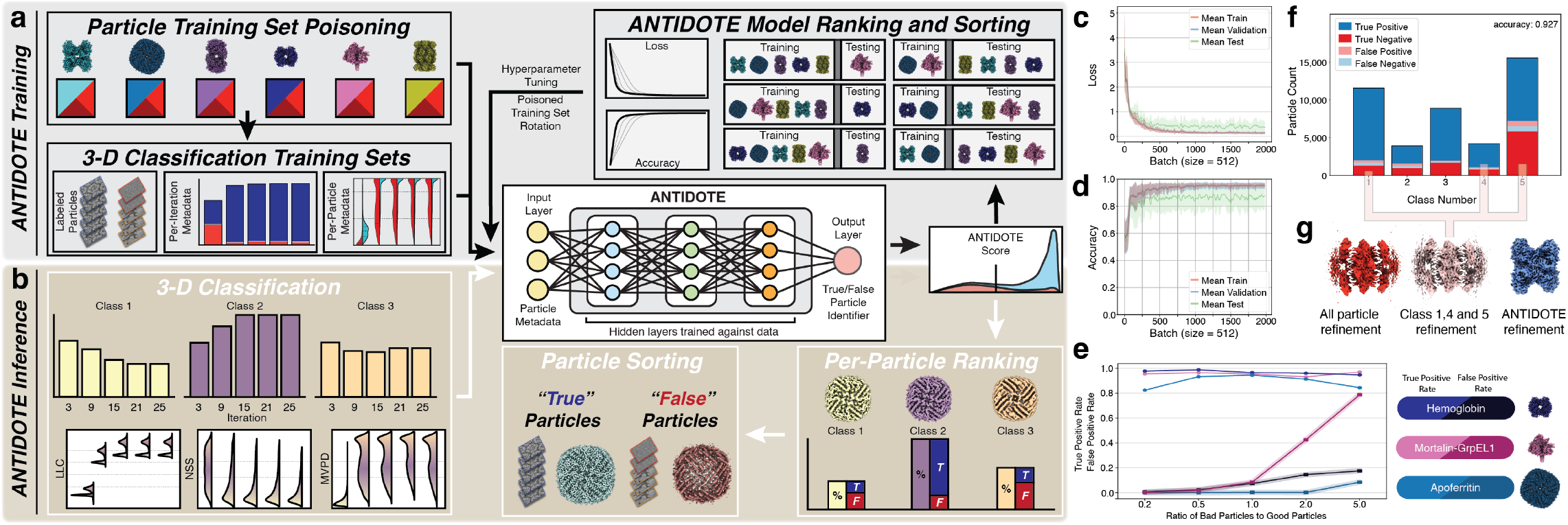
Ground-truth particle sets and expectation-maximization-based statistics from RELION 3-D classification leveraged to train ANTIDOTE. **a)** Schematic diagram of the ANTIDOTE training workflow. Ground-truth-labeled particle sets at varying True and False particle totals and True/False ratios were subjected to 3-D classification in RELION using an assortment of hyperparameters (*>*1000 total jobs). Ground-truth labels as well as per-iteration and per-particle metadata were used as inputs for generation of the ANTIDOTE binary classifier. A combination of training and testing specimen dataset ratios (e.g., 4 training and validation sets and 2 testing sets or 1-2 training and validation sets and 4-5 testing sets) were used for ANTIDOTE model ranking and sorting. This process was iterated through for final model generation. **b)** Schematic diagram of the ANTIDOTE inference workflow on real-world, unlabeled data. Metadata from a 3-D classification job in RELION is inputted into ANTIDOTE to generate per-particle confidence scores of 0 to 1 based on quality for sorting bad (False) from good (True) particles, respectively. **c)** Loss curves for ANTIDOTE training. **d)** Accuracy curves for ANTIDOTE training. **e)** Assessment of ANTIDOTE performance as a function of poison particle ratio. Specimens used for evaluation are shown. **f)** An example of particle classification using ANTIDOTE inference on an aldolase 3-D classification using the ground-truth-labeled evaluation dataset. **g)** 3-D reconstructions of the example aldolase data from (**f**) using all the particles in the initial particle stack (*left*; *red*), the optimal result from 3-D Classification (*middle*; *pink*), and after ANTIDOTE sorting (*right*; *blue*).

As a result of these efforts, we created ANTIDOTE: A Neural network Trained In Deleterious Object deTection and Elimination. ANTIDOTE is a binary particle classifier designed to operate solely on particle metadata and behaviors obtained during a RELION 3-D classification (**Fig. 3a, b**). Using metadata instead of the particle images themselves decouples ANTIDOTE’s performance from the molecular weight, dimensions, or nature of the biomacromolecule as well as the EM hardware used to acquire the data. Additionally, operating on features derived from 3-D classification instead of high-dimensional images allows for smaller, simpler models with fast inference times. The result of an ANTIDOTE analysis is a binary classifier that assigns every particle from a RELION 3-D classification job a confidence score between 0 and 1, corresponding to False and True, respectively, allowing the user to remove deleterious particles based on ANTIDOTE model classifications. The per-particle evaluation allows for a curation strategy that is independent of the particle’s class, and the use of a single neural-network-based approach that is robust to the parameters used in the input 3-D classification obviates the need for time-consuming parameter tuning currently involved in particle curation (**Fig. 3a, b**).

To prepare data for training or inference, ANTIDOTE employs a feature engineering approach that draws on meta-data contained in the data and model files generated by 3-D classification in RELION (see **Methods**). For each particle, ANTIDOTE parses the LLC, NSS, and MVPD values at each iteration of 3-D classification. In addition to these data, which are generated by default by RELION, ANTIDOTE applies a frequency encoding strategy to the class number and records the number of transitions between classes experienced by each particle over each iteration (**Fig. 3a, b**). Finally, ANTIDOTE adds target encoding features that map the class number for each particle to the Fourier completeness and estimated resolution of its assigned class. These features are drawn at each iteration to cumulatively describe the behavior of each particle during 3-D classification, totaling to a set of 241 features per particle from a standard 25 iteration RELION 3-D classification run. Each feature is normalized against the entire particle stack, providing context about the performance of each particle with respect to the particle stack (see **Methods**).

To identify the optimal model architecture, we evaluated a series of multilayer perceptron (MLP) architectures, along with MLPs prepended with a self-attention or multi-head self-attention module.^28–30^ Hyperparameter selection was performed using the hyperband approach and models were selected based on their validation accuracy.^31^ Final evaluation was conducted on an out-of-distribution test set containing proteins not seen during training or validation. In general, each combination of five of the six proteins in our training and validation set produced models that achieved high accuracy and low loss on the sixth protein, suggesting that there is an underlying signal in the statistical metadata from RELION 3-D classification and the engineered features generated by ANTIDOTE that can be leveraged for particle curation (**Fig. 3a-d**).

To evaluate the initial performance of the ANTIDOTE approach and to determine an optimal classification threshold, we trained models using the ANTIDOTE framework on two of the six proteins in our training dataset and cross-validated the resulting models against the 3-D classification jobs performed for the other four proteins (**Fig. 3a**). The two-protein models in our evaluation performed consistently well against the test protein datasets, with an average Youden’s J value of 0.80 and K-S statistic of 0.83 (p *<*0.001).^32^ In practice, to threshold the predictions generated by ANTIDOTE models, we provide two methods. By default, predictions are assigned a threshold of 0.5, a value that has been validated by analysis of the ROC curves generated by the dual-protein models described above. In addition, we provide an adaptive thresholder that returns particles in the rightmost cluster of the bimodal distribution of scores generated for most datasets by ANTIDOTE. Finally, we provide a utility alongside ANTIDOTE to choose a threshold value based on a histogram of predictions provided in a report generated alongside every ANTIDOTE inference run (**Fig. 3a, b**).

To illustrate ANTIDOTE’s potential for improving reconstruction quality through improved particle sorting, we ran inference on the poisoned aldolase dataset in **Fig. 2a** using a model trained on the other 5 proteins in our training dataset. ANTIDOTE recovers the high-quality aldolase particles and removes the ground-truth deconstructive particles regardless of their class (**Fig. 3f**), achieving a final accuracy of 92.7%. This curation results in an improvement of the aldolase reconstruction by 0.9 Å (**Fig. 3g**). The improvement in particle curation accuracy is seen across our panel of 3-D classification data when compared the accuracies achieved by 3-D classification-based particle curation alone (**Supplementary Fig. S7**)

### ANTIDOTE recovers particles mislabeled by standard classification approaches

Our initial classification evaluation efforts and subsequent ANTIDOTE training efforts included a T20S dataset commonly used as a benchmark for cryoEM methods development (EMPIAR-10025).^24^ Unlike the other five datasets in our panel, which are poisoned with False particles generated from micrographs that do not contain the protein of interest acquired on the same microscope, True and False labels for particles in the T20S datasets were generated using iterative rounds of 2-D classification in cryoSPARC using particles derived from the same over-picked micrographs (**Table 1**; see **Methods**). Particles in unambiguous 2-D classes containing high-quality T20S particles or obvious junk particles were assigned labels of True and False, respectively (**Fig. 4a, b**).

**Figure 4:**
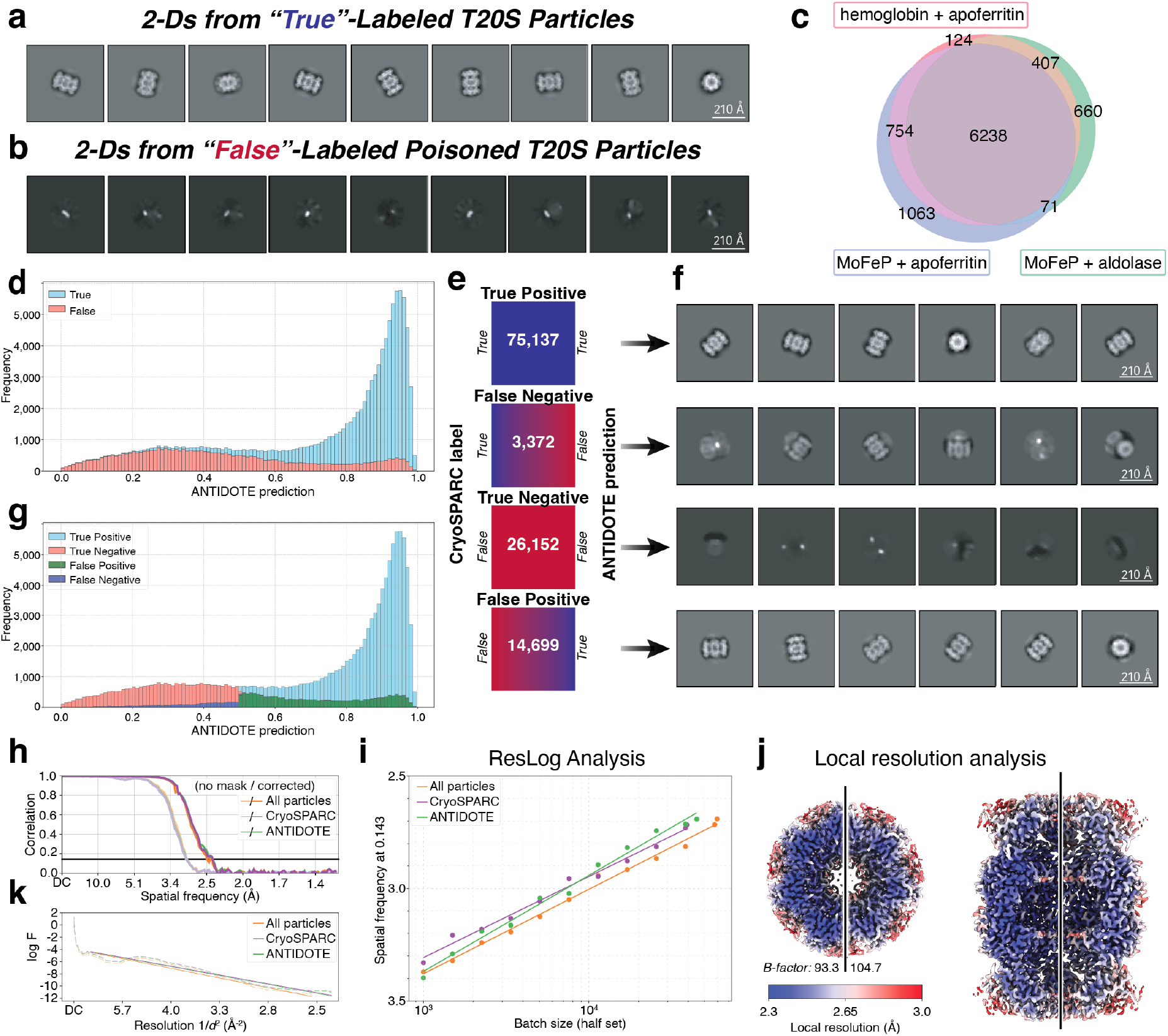
ANTIDOTE recovers particles incorrectly classified by 2-D classification. **a)** 2-D class averages for T20S proteasome particles initially labeled as “True” using curation in cryoSPARC. **b)** 2-D class averages for T20S proteasome particles initially labeled as “False” using curation in cryoSPARC. **c)** A Venn diagram for False positives identified by ANTIDOTE models trained on orthogonal sets of training data. **d)** The distribution of ANTIDOTE predictions for the T20S proteasome dataset with ground-truth-labeled particles colored by True (blue) or False (red) labels. **e-f)** Confusion matrix for ANTIDOTE predictions using the original cryoSPARC labels as the ground-truth and representative 2-D class averages of the particles shown in (**f**). **g)** The distribution of ANTIDOTE predictions for the same T20S proteasome data in d-f) with updated ground-truth-labels after one round of ANTIDOTE inference. **h-k)** Fourier shell correlation curves (**h**), Guinier plots (**i**), and ResLog plots (**k**) of the full, cryoSPARC-derived, and ANTIDOTE-derived particle stacks, and local resolution estimates for the T20S proteasome reconstructions generated using ANTIDOTE particles (*left*) and cryoSPARC particles (*right*) (**k**).

Interestingly, initial tests of ANTIDOTE models with this dataset led to unexpectedly low maximum accuracies of around 80%. To investigate this discrepancy more thoroughly and mitigate potential biases from any single set of 3-D classification parameters, we performed ANTIDOTE inference on all 108 T20S 3-D classification jobs within our evaluation dataset, each employing a diverse range of parameters, and then calculated a mean ANTIDOTE score for each particle across all the classification jobs (**Fig. 4d**). While the distribution of ANTIDOTE predictions in this dataset was bimodal, a significant population of false positives is present, including a small peak of False particles in the high ranges of the ANTIDOTE prediction histogram (**Fig. 4d**). This result suggests that either ANTIDOTE is misclassifying deconstructive particles as True, or that the original labeling performed by 2-D classification in cryoSPARC misclassified a subset of constructive particles as False.

To evaluate this discrepancy and determine whether ANTIDOTE is correctly identifying particles that were incorrectly labeled by cryoSPARC 2-D classification (as opposed to True false positives that are mislabeled by ANTIDOTE), we compared particle predictions for ANTIDOTE models trained on two proteins from our panel of training data in order to compare models trained on orthogonal sets of our training data. We found significant overlap in the particles labeled as false positives by two-protein ANTIDOTE models — across three of the best two-protein ANTIDOTE models trained on orthogonal data, we found that 67% of the false positives identified across the three models were unanimous across the three models (**Fig. 4c**). We also trained a CryoDRGN model and found that a significant portion of the particles in the latent space cluster corresponding to constructive T20S particles were labeled as False (**Supplementary Fig. S8**).^33^ We also found significant correlation with ANTIDOTE labels and the latent space clusters generated by CryoDRGN (**Supplementary Fig. S8**).

Analysis of each quadrant of the confusion matrix by 2-D classification in cryoSPARC provides further confirmation that ANTIDOTE is correctly identifying constructive and deconstructive T20S particles that are incorrectly classified by 2-D classification of the full particle stack in cryoSPARC (**Fig. 4e-g**). As expected, 2-D classification of true positives (particles included in good classes by cryoSPARC and given labels above 0.5 by ANTIDOTE) generated high-quality 2D class averages with clear top and side views of T20S, and true negatives generated class averages with poor contrast that appear to be driven by noise rather than protein signal (**Fig. 4e, f**). Interestingly, 2-D classification of the false positives (particles included in low-quality 2-D classes by cryoSPARC but assigned values greater than 0.5 by ANTIDOTE) generates a significant number of high-quality classes (**Fig. 4e, f**). This suggests that a significant population of constructive particles were rejected during initial 2-D classification-based curation of the full particle stack, and that a sufficient population of these particles are recovered by ANTIDOTE to generate high-quality 2-D class averages. Finally, 2-D classification of false negatives (particles labeled as True in the initial curation campaign but labeled as False by ANTIDOTE) generates 2-D class averages containing misaligned or crowded particles (**Fig. 4e, f**). Despite containing T20S particles, these particles would likely be deconstructive in the context of a high-resolution reconstruction of T20S and are therefore appropriately rejected by ANTIDOTE despite originally being included in high-quality 2-D class averages by 2-D classification in cryoSPARC. An updated histogram of ANTIDOTE predictions reflects the original 2-D classification labeling errors and aligns with the downstream confusion matrix 2-D classifications, revealing instances where particles that should likely be ground truth negative are labeled as ground truth positive (below the 0.5 threshold) and, conversely, particles that should likely be ground truth positive are labeled as ground truth negative (above the 0.5 threshold) (**Fig. 4g**).

While the results of relabeling the T20S dataset using ANTIDOTE is most evident in the 2-D classes of particles in each quadrant of the label confusion matrix (**Fig. 4e, f**), reconstruction of the ANTIDOTE- and cryoSPARC-derived particle stacks reveals a marginal improvement in the nominal FSC-estimated resolution of the T20S volume (**Fig. 4h-k**). Despite similar bulk reconstruction metrics provided by ResLog, GSFSC, and Guinier plots for both the ANTIDOTE- and cryoSPARC-derived reconstructions as well as reconstructions derived from the full particle stack (**Fig. 4h-k**),^11,34,35^ an analysis of the local resolution of the ANTIDOTE- and cryoSPARC-derived reconstruction shows that correcting the labeling errors derived from the original 2-D classification can yield an improvement reconstruction quality even when dealing with an inherently robust dataset like the T20S proteasome (**Fig. 4j**).

### ANTIDOTE accelerates the generation of high-quality 3-D reconstructions with minimal manual particle curation

We initially evaluated ANTIDOTE’s performance on real-world data using a hemoglobin dataset recently published by our group (EMDB-45816; EMPIAR-12743). Hemoglobin (Hb) is a 64 kDa heterotetramer that is a particularly challenging target for particle curation due to its small size. Initial reconstruction of the Hb volume required significant manual particle curation.^23^ To evaluate ANTIDOTE’s performance on datasets with varying degrees of manual curation, and to provide insight into integration approaches for ANTIDOTE into the standard cryoEM data processing workflow, we prepared three Hb particle stacks: an uncurated stack, a partially curated stack, and a curated stack. Each particle stack is designed to evaluate the performance of ANTIDOTE at different stages of a real-world particle curation campaign (**Fig. 5**).

**Figure 5:**
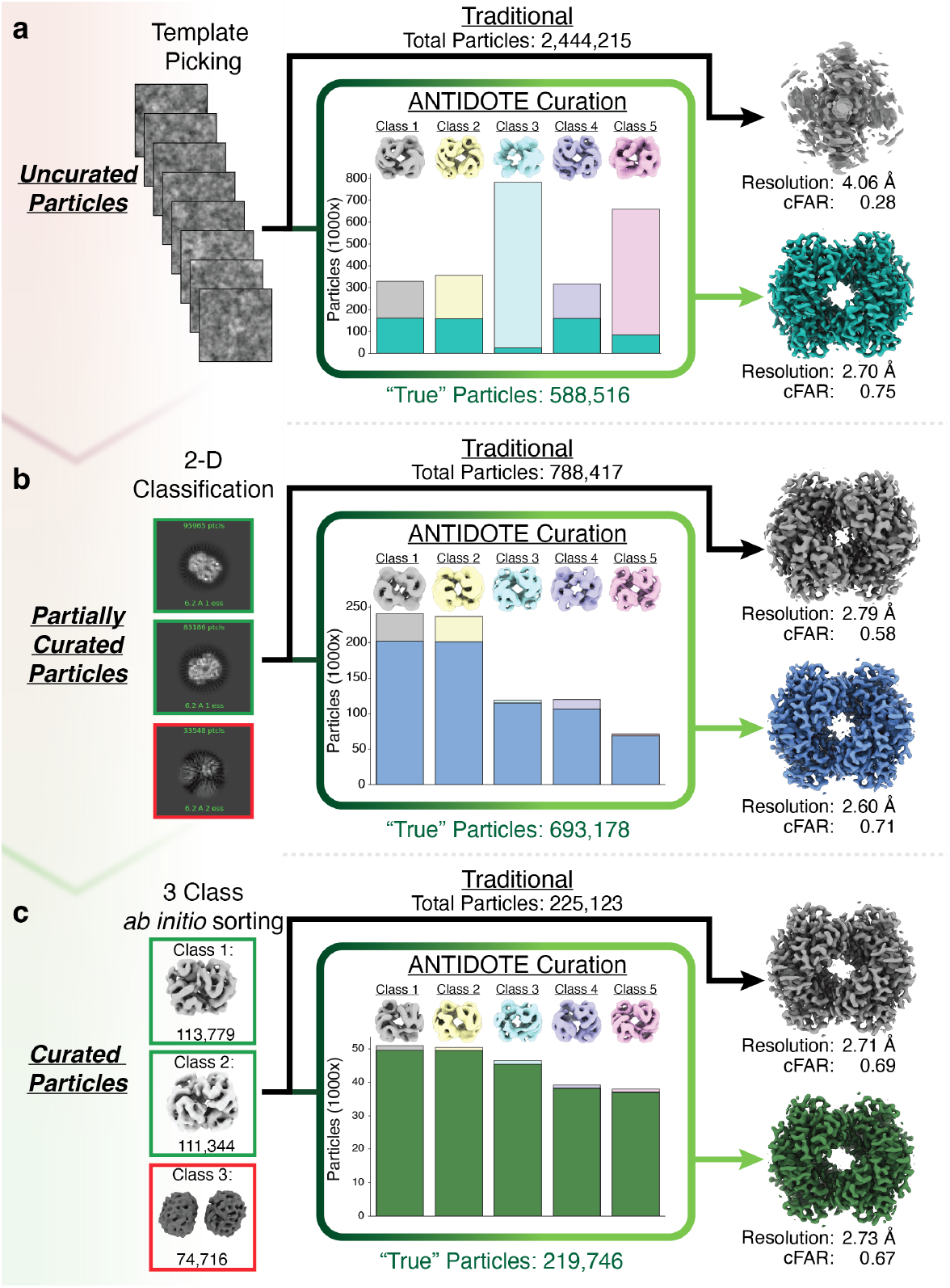
ANTIDOTE inference on minimally-curated particle stacks can outperform traditional classification approaches. **a)** Hemoglobin particle images obtained from template-based particle picking (*left*). These uncurated particles (2,444,215) were either directly refined (gray volume; *right*) or subjected to 3-D classification (K=5; T=4) followed by ANTIDOTE inference (588,516 particles) and refinement (teal volume; *right*). **b)** The uncurated particles from (**a**) were subjected to 2-D classification. 2-D class averages (*left*) with secondary structure features (green box; 788,417 particles) were either directly refined (gray volume; *right*) or subjected to 3-D classification (K=5; T=4) followed by ANTIDOTE inference (693,178 particles) and refinement (blue volume; *right*). **c)** The partially curated particle stack from (**b**) was subjected to a 3-class ab initio and the best volumes (green box; 225,123 particles) were either directly refined (gray volume; *right*) or subjected to 3-D classification (K=5; T=4) followed by ANTIDOTE inference (219,746 particles) and refinement (green volume; *right*).

The first particle stack contained template-picked particles that had not been curated by 2-D classification. *Ab initio* reconstruction and refinement using the full, uncurated stack of 2,444,215 particles resulted in an uninterpretable, noisy volume with a conical FSC Area Ratio (cFAR) of 0.28 (**Fig. 5a**). The sensitivity of the cFAR metric to particle quality suggests that although this dataset could have significant orientational bias, it is more likely that the uncurated particle stack consists largely of noise and deleterious particles. Despite this significant percentage of contaminating deconstructive particles, running ANTIDOTE inference on 3-D classification results from the uncurated particle stack removed nearly 75% of the particles from the initial stack (*∼*2.44M to *∼*589K particles) to yield a more interpretable 2.70 Å reconstruction with a cFAR of 0.75 (**Fig. 5a**). The performance of ANTIDOTE on this completely uncurated particle stack suggests that intervention-free processing with a single round of 3-D classification using the default parameters followed by ANTIDOTE inference could be a viable curation strategy (**Fig. 5a**), although the significant time investment required for 3-D classification of such a large particle stack makes it unlikely that this would be a reasonable first approach to take.

Next, the initial Hb particle stack generated from particle picking was subjected to a single round of 2-D classification to generate a partially curated Hb particle stack of 788,417 particles. *Ab initio* reconstruction and refinement of this partially curated particle stack without ANTIDOTE classification generates a Hb volume with an estimated resolution of 2.79 Å and a cFAR of 0.58 (**Fig. 5b**). In comparison, running ANTIDOTE inference on the partially curated particle stack produces a reconstruction with a resolution of 2.60 Å and a cFAR of 0.71. This improvement in resolution and cFAR is attained despite the removal of a modest 12% of the particle stack (**Fig. 5b**). ResLog analysis of the partially curated Hb particle stack with and without ANTIDOTE curation confirms that the particles selected by ANTIDOTE consistently generate higher-resolution reconstructions than those derived from partial curation alone (**Supplementary Fig. S9**).

Finally, to represent a fully curated Hb particle stack we performed a 3-class *ab initio* reconstruction on the partially curated particle stack. We then selected the two best classes (highlighted in green) based on volume completeness and used these particles to represent the curated Hb particle stack (**Fig. 5c**). After classification in 2-D and 3-D, few deleterious particles remained in the curated particle stack. Accordingly, ANTIDOTE inference removed only *∼*2% of the particle stack, producing a particle stack whose reconstruction is similar in resolution to refinement of the curated particle stack (**Fig. 5c**). ResLog analysis of the curated particle stack and the ANTIDOTE-treated curated particle stack reveals near-identical slopes, implying that the particle stacks are nearly identical in their quality (**Supplementary Fig. S9**). While ANTIDOTE’s retention of constructive particles in clean particle stacks suggests that it is a balanced classifier even in the absence of deconstructive particles, its use after extensive manual curation may often be superfluous.

### Improvement of real-world cryoEM datasets using ANTIDOTE

In addition to demonstrating ANTIDOTE’s ability to enhance cryoEM data processing workflows by reducing the amount of practitioner input necessary to achieve a high-resolution reconstruction, we chose to further validate ANTIDOTE’s ability to generalize and to improve the quality of a reconstruction by applying ANTIDOTE to three published cryoEM datasets: the nitrogenase molybdenum-iron protein (MoFeP, EMPIAR-48384), the bacterial chaperonin GroEL (EMPIAR-11304), and the human calcium homeostasis modulator 1 channel (CALHM1, EMPIAR-11668; **Fig. 6**).^23,36,37^ These datasets represent diverse molecular weights, symmetries, and data collection parameters that allow us to assess ANTIDOTE’s performance across a range of experimental conditions.

**Figure 6:**
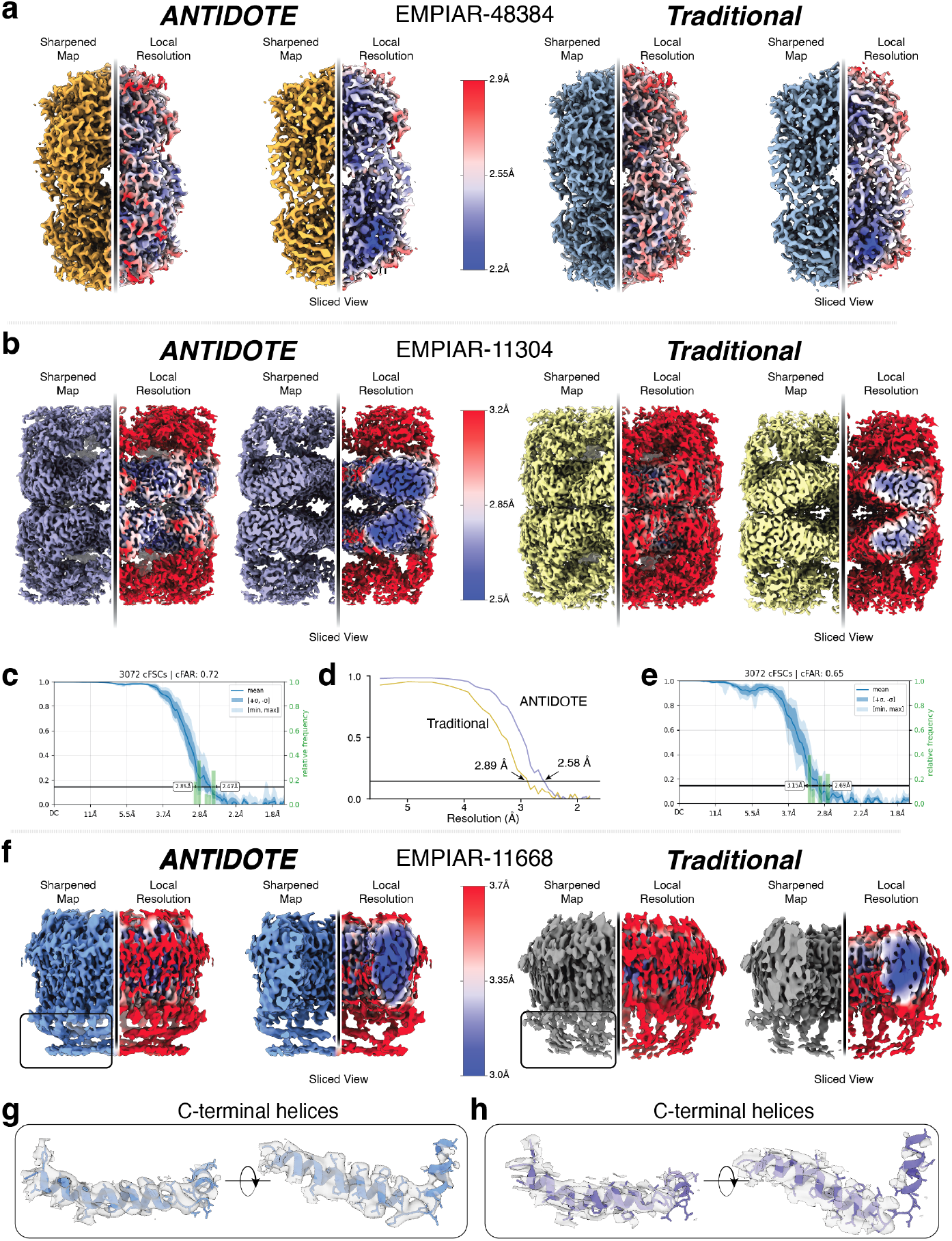
ANTIDOTE matches or improves upon traditional classification approaches. Comparison of 3-D reconstruction quality following ANTIDOTE inference (*left*) compared to Traditional processing (*right*) for EMPIAR-48384 (**a**), EMPIAR-11304 (**b-e**), and EMPIAR-11668 (**f-h**). The FSC curves indicating resolution improvements from ANTIDOTE for (**b**) are shown in panels (**c-e**). **g-h)** The C-terminal helices from (**f**) are shown to indicate map quality improvement from ANTIDOTE.

We first evaluated ANTIDOTE’s performance on MoFeP, a C_2_-symmetrical heterotetrameric metalloprotein that is relatively small as compared to CALHM1 and GroEL. Micrographs containing MoFeP were blob-picked, subjected to one round of 2-D classification, and exported to RELION for 3-D classification followed by ANTIDOTE inference (**Supplementary Fig. S10**; see **Methods**). The particles identified by ANTIDOTE as being constructive were then 3-D refined using cryoSPARC. For comparison, the particles selected by 2-D classification were similarly refined in cryoSPARC. ANTIDOTE was able to discard *∼*20K deleterious particles from the original particle stack and achieve a moderate resolution gain of 2.41 Å for ANTIDOTE as opposed to 2.49 Å for the original particle stack (**Fig. 6a**). This increase in resolution while using less particles shows that during standard curations there may be either deleterious or at least non-contributing particles remaining within the particle stack, regardless of curation strategies.

We next sought to understand ANTIDOTE’s performance on a large protein with high symmetry that has been used as a common benchmark sample for the evaluation of cryoEM methods. To this end, we selected GroEL, a large protein complex that comprises 14 protomers and exhibits D_7_ symmetry. We used a simple blob picker to generate a particle stack which was then subjected directly to 3-D classification in RELION followed by ANTIDOTE inference (**Supplementary Fig. S11**). Interestingly, while RELION sorts a significant set of GroEL particles into a single class, ANTIDOTE selects only a subset of the major RELION class while also retaining most of the particles in the four additional minor classes (**Supplementary Fig. S11**). For the classical curation pipeline, we selected the two best classes from the 3-D classification for comparison with the ANTIDOTE results. Both the ANTIDOTE and the classical curation particle stacks were imported into cryoSPARC and refined. Reconstruction of the ANTIDOTE particle stack yields a 2.58 Å reconstruction with a cFAR of 0.72, outperforming the classical approach, which reaches a resolution of 2.89 Å with a cFAR of 0.65 (**Fig. 6b-e**). The local resolution of the GroEL reconstruction is markedly improved in the ANTIDOTE density compared to the density generated by classical curation alone, possibly due to the improvement in viewing angle distribution evidenced by the high cFAR of the ANTIDOTE-derived reconstruction (**Fig. 6c,e**). The improvement in viewing angle distribution and reconstruction resolution afforded by ANTIDOTE illustrates its potential to directly improve data reconstruction by drawing subsets of particles directly from 3-D classification classes, especially in cases where RELION sorts particles into a single major class (**Fig. 6c-e**).

To complete our evaluation of a diverse set of biomacromolecules, we examined ANTIDOTE’s performance on the octameric voltage-gated ion channel CALHM1. CALHM1 is an integral membrane protein that is solubilized in nanodiscs for single-particle cryoEM data acquisition. CALHM1 has an extensive C-terminal helix domain on the cytoplasmic side of the membrane that exhibits weaker density compared to the transmembrane domains and appears to be susceptible to oversharpening.

To evaluate ANTIDOTE performance on this membrane protein, the data were repicked using a simple blob picker and subjected to 3-D classification in RELION for ANTIDOTE inference. From this 3-D classification, we generated two particle stacks, an ANTIDOTE-selected and the best class from the 3-D classification. Both particle stacks were then subjected to a 5-class *ab initio* and the volumes that resembled CALHM1 were selected (**Fig. 6f, Supplementary Fig. S11**; see **Methods**). Despite the novelty of the CALHM1 data compared to our training data and the distribution of particles across 5 classes by RELION, ANTIDOTE was able to identify particles in each class that contributed positively to a single homogeneous final map (**Fig. 6f, Supplementary Fig. S11**). While the global resolution of both reconstructions is nearly identical, closer inspection of the maps shows that the ANTIDOTE reconstruction has interpretable density representing the C-terminal basket-like helices whereas the map resulting from the standard curation approach has only weak and discontinuous density in this region (**Fig. 6f-h**). This additional density in the ANTIDOTE-derived map supports model building in the C-terminal helices that could not be performed in the classically-prepared map (**Fig. 6f-h**). Additionally, the local resolution of the two maps shows that the core of each protomer in the ANTIDOTE map exhibits higher resolution and more interpretable density (**Fig. 6f**). Overall, while ANTIDOTE may have not increased the global resolution, its application results in a higher quality and more complete density that supports additional atomic model building.

## Discussion

The reconstruction of high-resolution macromolecular structures using cryoEM relies on the effective sorting and curation of particle images that constructively contribute to the final 3-D volume. Curation is challenging due to the low signal-to-noise inherent in cryoEM data and the lack of *a priori* information about the ground-truth system represented in the particle stack. Modern approaches rely on Bayesian methods that attempt to simultaneously solve the particle classification problem and determine the 3-D structure of the macromolecule represented in the data.^8,13^ While this approach is incredibly effective, the reliance on iterative classification into a small number of 2-D and 3-D classes leads to the inevitable loss of constructive data, especially early in the process when uncertainty about the macromolecular structure is highest and the proportion of deconstructive particles in the dataset is large.^7^ This particle misclassification represents a significant opportunity cost, especially for challenging datasets where particle numbers are limited.^7^ Additionally, current methods involve exploratory hyperparameter tuning that can consume a significant amount of time, even for expert practitioners.

In this study, we generated a broad set of ground-truth-labeled particle sets by treating heavily curated data as ground-truth True particles and junk particles produced from separate data acquired on the same EM instrumentation as ground-truth False particles. We then proceeded to characterize the performance of standard 2-D and 3-D data processing approaches to quantify the opportunity cost inherent in current processing workflows (**Fig. 1-2**). Importantly, our analysis reveals that the ratio of True-to-False particles has a large impact on the retention of constructive data and the exclusion of destructive data — when a large proportion of the particle population is False, many True particles are incorrectly binned into False classes, especially during 2-D classification, which is more frequently run early in data processing workflows when the proportion of False particles is large (**Fig. 1e, f**). Similarly, we find that when more True particles are present in the data, real differences in features in the data can drive particle classification boundaries, in both 2-D and 3-D, leading to the distribution of False particles among classes that are primarily differentiated by features in the True particles (**Fig. 1h-j; Fig. 2a c, f,; Fig. 3e, f**). While these results may be expected due to the nature of the classification algorithms used in particle curation, they highlight an opportunity for more effective particle classification approaches that operate on a per-particle basis.

In addition to evaluating 2-D and 3-D classification, we leverage our labeled data to train ANTIDOTE, a neuralnetwork-based binary classifier framework that ameliorates the opportunity cost represented by misclassified particles by accurately assigning True and False labels to both internal and external datasets (**Fig. 3a, b**).^38^ ANTIDOTE ingests the per-particle Bayesian statistics generated by 3-D classification and combines these statistics with additional information from RELION reporting on particle and volume quality to construct a dataset that represents each particle’s behavior across each iteration of 3-D classification. ANTIDOTE then passes the normalized data for each particle through a multilayer perceptron trained on hundreds of 3-D classifications of our ground-truth labeled particles to generate a classification for each particle. ANTIDOTE therefore leverages the rich set of statistics generated during RELION 3-D classification to produce an effective discriminator of particle quality (**Fig. 3a**).

Our extensive evaluation of particle classification with ANTIDOTE demonstrates that this approach often outperforms 2-D or 3-D classification on a given particle stack, especially at early stages of particle curation when deconstructive particles are abundant (**Fig. 3e-g, Fig. 4, Fig. 5, Fig. 6**). The effectiveness of ANTIDOTE (and the RELION 3-D classification algorithm that supports it) on diverse datasets, spanning a range of molecular weights, symmetries, and contamination levels, demonstrates the robustness of our approach and its potential to generalize to new datasets. Additionally, widespread use of 3-D classification and the short inference time of ANTIDOTE (seconds to minutes) means that cryoEM practitioners can install and rapidly apply ANTIDOTE to preexisting 3-D classification data. The ability of ANTIDOTE to outperform standard data processing workflows while also reducing the time and manual involvement in data processing workflows position it as a widely applicable tool for cryoEM data processing. Importantly, ANTIDOTE can recover constructive particles that would otherwise be discarded during conventional classification approaches while simultaneously removing deconstructive particles that might be retained, resulting in higher-quality reconstructions with improved resolution and more uniform local resolution distributions.

Based on our observations, we recommend incorporating ANTIDOTE early in the particle curation pipeline, using downsampled data that have ideally through an initial round of 2-D classification or the generation of a reasonable 3-D structure with discernable features. While ANTIDOTE can process completely uncurated datasets, its performance is improved when some structural features begin to appear in the classes generated by RELION (**Fig. 5a,b**). It is worth noting that significant computational time can be required for 3-D classification of uncurated data, so a minimal initial curation step using 2-D classification to conservatively remove obvious deconstructive particles is often worthwhile. Importantly, ANTIDOTE is designed to enhance the reconstruction quality of homogeneous datasets and therefore serves as an orthogonal and complementary tool to recent advances in handling heterogeneity in cryoEM data.^33,39^

As cryoEM continues to evolve as a primary method for the structural determination of biological macromolecules, automation in the data processing pipeline will allow practitioners to focus on generating and interpreting new data rather than iteratively adjusting established data processing parameters. The synergy of ANTIDOTE with RELION’s framework, coupled with its rapid inference performance, often on the order of seconds for standard datasets, means that ANTIDOTE can promptly be applied concurrently with 3-D classification. Indeed, we show that ANTIDOTE improves upon traditional particle class assignments generated during 3-D classification by accessing statistics that describe each particle’s behavior at each step of the iterative 3-D classification process. This wealth of information enables ANTIDOTE to discriminate between constructive and destructive particles, irrespective of their final 3-D class assignment, increasing the accuracy of 3-D classification, and ultimately, the quality of high-resolution 3-D reconstruction. ANTIDOTE therefore allows even a novice user to quickly remove deconstructive, or “poisonous” particles from their data without the time-consuming hyperparameter scouting approaches and subjective decision making that are currently required.

In conclusion, we show that ANTIDOTE can serve as a key component in this automation by decreasing the impact of subjective decision-making in the cryoEM data processing workflow. ANTIDOTE represents a valuable addition to the cryoEM data curation toolbox, serving two key purposes. First, it streamlines parameter by replacing time-consuming manual experimentation with automated particle curation pipelines. Second, ANTIDOTE can extract additional resolution and map quality from homogeneous datasets by achieving more accurate particle curation than conventional class-based approaches. By combining the effectiveness of established Bayesian classification methods with the pattern recognition capabilities of neural networks, ANTIDOTE provides a robust, user-friendly approach that advances the field of cryoEM structure determination toward more consistent, high-quality results with reduced user intervention.

## Methods

### Generation of “True”-labeled particle datasets

#### Rabbit muscle aldolase

Particles were obtained from the final particle stack used to generate EMD-21023 (0.56 Å/pixel).^20^ These particles (179,950) were clipped to a box size of 384 pixels and then down sampled (96-pixel box, 2.26 Å/pixel) for use as a “True” particle set (**Supplementary Fig. S1**).

#### Mouse heavy chain apoferritin

Particles were obtained from the final stack used to generate EMD-21024 (0.566 Å/pixel).^20^ These particles (241,878) were down sampled (96-pixel box, 2.264 Å/pixel) for use as a “True” particle set (**Supplementary Fig. S1**).

#### Human Hemoglobin (Hb)

Initial particle coordinates were acquired from the original particle picks that were ultimately used to generate EMD-45815 (raw data located at EMPIAR-12743).^23^ Briefly, these initial picks (2,444,215) were obtained from template-based particle picking using EMD-45815 and extracted using a box size of 256 pixels and Fourier cropped to 64 pixels (2.94 Å/pixel). These particles were subjected to two iterative rounds of 2-D classification followed by *ab initio* model generation (3 classes, C_1_ symmetry). The class most resembling Hb was selected for non-uniform refinement (C_2_ symmetry) to yield a 6.38 Å resolution reconstruction. This particle stack was re-extracted with a box size of 352 pixels (0.735 Å/pixel) and subjected to a non-uniform refinement (C_2_ symmetry) with CTF and aberration refinements. To further clean these particles, the particle stack was reextracted (64-pixel box, 2.94 Å/pixel) and subjected to four rounds of iterative 2-D classification. Only classes displaying strong Hb features were selected and these particles (241,409) were exported to RELION-5.0 using csparc2star from pyem for use as a “True” particle set.^18,40,41^ To ensure these particles would achieve high resolution, this final particle stack was re-extracted (352-pixel box, 0.735 Å/pixel) and a final non-uniform refinement (C_2_ symmetry) with CTF and aberration refinements to yield a 2.46 Å resolution reconstruction (**Supplementary Fig. S2**).

#### Human mortalin-GrpEL1 complex

A final particle stack of 51,253 human mortalin-GrpEL1 complex particles (obtained from EMD-44675, 0.735 Å/pixel)^22^ that yielded a 3.1 Å resolution reconstruction using non-uniform refinement were re-extracted with a 96-pixel box (2.205 Å/pixel) and exported to RELION-5.0 using cspar2star from pyem for use as a “True” particle set (**Supplementary Fig. S2**).^18,40,41^

#### *Azotobacter vinellandii* nitrogenase molybdenum-iron protein (MoFeP)

Particles were initially selected from the aligned micrographs that were used to obtain EMD-26757 using a template-based particle picker with EMD-26757 as a reference.^21^ These particles (3,064,350) were then subjected to two iterative rounds of 2-D classification resulting in 728,079 particles. These particles were then recentered and re-extracted (64-pixel box, 3.416 Å/pixel) and exported to RELION-5.0 using csparc2star from pyem for use as a “True” particle set.^18,40,41^ These particles were then re-extracted at a box size of 384 pixels with a pixel size of 0.854 Å/pixel and a final non-uniform refinement (C_2_ symmetry) with CTF and aberration refinements resulting in a 2.27 Å resolution reconstruction (**Supplementary Fig. S3**).

#### *Thermophilus acidophilum* 20S proteasome core (T20S)

Raw micrographs from EMPIAR 10025 (196 movies; 7420 x 7676 pixels, 0.66 Å/pixel, 40 electrons/Å^2^) were patch motion corrected (Fourier cropped to 1.32 Å/pixel) and patch CTF estimated within cryoSPARC v4 6.2.^9,24,42,43^ Particles were selected via a blob picker and extracted with a box size of 384 pixels (1.32 Å/pixel) with Fourier cropping to 96 pixels (5.28 Å/pixel). Extracted particles (371,759) were then subjected to four iterative rounds of two-dimensional (2-D) classification where 2-D class averages containing strong T20S signal were selected for downstream processing (126,737 particles). *Ab initio* model generation (2-classes, C_1_ symmetry) yielded one prominent T20S class (80,632 particles) that was selected for a non-uniform refinement (C_1_ symmetry).^9^ These particles were then extracted without binning (384-pixel box, 1.32 Å/pixel) and subjected to reference-based motion correction. A final non-uniform refinement with CTF and aberration refinements yielded a 2.09 Å resolution reconstruction (D_7_ symmetry).^42^ The resulting 78,509 particles were then re-extracted with a box size of 128 pixels (1.98 Å/pixel) and were then exported to RELION-5.0 using csparc2star from pyem for use as a “True” particle set.^18,40,41^

### Generation of “False”-labeled poison datasets

#### Aldolase and apoferritin

To generate “False” particle stacks for poisoning aldolase and apoferritin, images of a crossed-lines grating replica calibration grid (19 movies; termed “calibration”) or over vacuum (11 movies; termed “noise”), respectively, were recorded using a Talos Arctica (ThermoFisher Scientific) operating at 200 kV equipped with a Gatan K2 Summit Direct Electron Detector (DED). Movies were collected at a nominal magnification of 73,000x in counting mode with a cumulative electron exposure of 66 electrons/Å^2^. These parameters were chosen to match those used for aldolase (EMD-21023) and apoferritin (EMD-21024) which were all collected at the CryoEM facility at Scripps Research using the identical instrument and data collection parameters.**1** Data were collected using Leginon.^44^ Micrographs were motion corrected using motionCor2 within RELION-3.0v2 and CTF estimation was performed using GCTF.^41,45,46^ Particle coordinates were obtained by generating a matrix of picks across each micrograph equally spaced in X and Y dimensions yielding 12,125 particles per micrograph. 2-D “poison” images were then extracted at a 384-pixel box size (Fourier cropped to 96 pixels) and scaled to match the “True” particle set (aldolase, 2.26 Å/pixel; apoferritin, 2.264 Å/pixel). Each particle set was then exported to RELION-3.0v2 using csparc2star from pyem for combining with the aldolase or apoferritin “True” particle sets in varying ratios (**Supplementary Fig. S1**).

#### Hb and mortalin-GrpEL1

Contaminant poison particles used in the Hb and mortalin-GrpEL1 datasets were obtained from movies collected on manually-frozen cryoEM grids using water from a 37 °;C water bath. Briefly, 3 *µ*L of “water” was applied to a UltraFoil R1.2/1.3 grid that had been glow discharged using a Gatan Solaris II plasma cleaner (10 s, 15 Watts, 75% Ar/25% O_2_ atmosphere) and manually blotted using a custom manually-operated plunging device for 6-8 s with Whatman #1 blotting paper.^47^ 446 movies were collected using a Titan Krios G4 (ThermoFisher Scientific) operating at 300 kV equipped with a Selectris-X energy filter (ThermoFisher Scientific) with a nominal magnification of 165,000x in EF-TEM mode (with a calibrated pixel size of 0.735 Å) on a Falcon 4 DED (ThermoFisher Scientific) using a 10 eV slit width. Movies were acquired in EER format with a cumulative electron exposure of 60 electrons/Å^2^ with a defocus range of −1 to −2.0 *µ*m. These parameters were chosen to match those used for Hb (EMD-45815) and mortalin-GrpEL1 (EMD-44675) which were all collected at the CryoEM facility at UCSD using identical instrument and data collection parameters.^22,23^ Data were collected automatically using EPU2 (ThermoFisher Scientific) using aberration free image shifting. Movies were patch-based motion corrected and patch CTF estimated in cryoSPARC v4 6.2. Particle stacks were generated by using a template-based picker using a Hb volume as a template (20 Å low-pass, 90 Å particle diameter, 30 Å minimum spacing) without any further curation. This resulted in 1,365,388 picks that were subsequently extracted at the required box size to match the “True” particle dataset (256-pixel box, 2.94 Å/pixel for Hb; 384-pixel box, 2.205 Å/pixel for mortalin-GrpEL1). Each particle set was then exported to RELION-5.0 using csparc2star from pyem for combining with the Hb or mortalin-GrpEL1 “True” particle sets in varying ratios (**Supplementary Fig. S2**).^18,40,41^

#### MoFeP

Particles for MoFeP were generated using the deselected particles from the first round of 2-D classification used to generate the “True” MoFeP particle stack. These contaminant particles (1,964,704) were subjected to a second round of 2-D classification and any classes containing discernable protein features were discarded. This was repeated an additional time until no obvious protein classes were present to yield a final particle stack contained 553,441 particles. These particles were then re-extracted (64-pixel box, 3.416 Å/pixel) and exported to RELION-5.0 using csparc2star from pyem for combining with the MoFeP “True” particle sets in varying ratios (**Supplementary Fig. S3**).^18,40,41^

#### T20S proteasome core

Poison particles for T20S were generated by using the de-selected particles from the first round of 2-D classification that was used to generate “True” T20S particles. These contaminant particles (60,522) were then subjected to a second round of 2-D classification and any 2-D classes resembling T20S were discarded. This was repeated an additional time until no obvious protein classes were present yielding a total of 40,857 “False”-labeled poison particles. These particles were then re-extracted with a 128-pixel box (1.98 Å/pixel) and then exported to RELION-5.0 using csparc2star from pyem for combining with the T20S “True” particle sets in varying ratios (**Supplementary Fig. S3**).^18,40,41^

### Generating poisoned particle stacks for 3-D classification

#### Aldolase, apoferritin, Hb, MoFeP, mortalin-GrpEL1, and T20S

To generate STAR files for training within RELION, each labeled data set used the following procedure. “True” particle sets were split randomly into different subsets ranging from 10% of the total data to the full particle stack (see **Table 1**). Poison particles were then randomly selected at different percentages, ranging from 5% to 50% of the “True” particle stack. These particle stacks were merged to form a final STAR file which was subsequently used for generating 3-D classification training data. For example, 50,000 randomly-selected “True” Hb particles (out of a total 241,409 available) would be mixed with 5% (2,500 particles) random contaminant poison particles (out of a total of 1,365,388 particles). This mixing was then done iteratively for each desired “True” particle amount and ratio of contaminant “False” poison particles. Note, for each combination, a new random selection for both “True” and “False” particles was performed to limit the extent of overlap and/or the repeated use of the same stacks for training purposes. This is summarized in SI **Supplementary Fig. S1-3**.

### 3-D classification to generate training data

Training datasets were generated by using 3-D classification within RELION-3.0 for apoferritin and aldolase.^41^ Training datasets were generated by using 3-D classification within RELION-5.0 for T20S, Hb, Mortalin-GrpEL1 and MoFeP.^48^ Class number (e.g., 2, 3, or 5) and T regularization values (e.g., 1, 3, or 7) were simultaneously varied across datasets to create a robust training set (1074 total 3-D classification; see **Table 1**). These values were selected to mimic the different 3-D classification parameters a user might run during processing. Each combination of K and T values were run on previously generated “poisoned particle” stacks. The following training sets were generated:

#### Aldolase

A total of 54 3-D classification jobs using a combination of 2, 3, or 5 classes with varying T values of 1, 2, and 4 and E-steps of −1 or 6 for each particle amount and poison percentage (**Supplementary Fig. S1**; **Table 1**).

#### Apoferritin

A total of 192 3-D classification jobs using a combination of 2, 3, or 5 classes with varying T values of 1, 2, and 4 and E-steps of −1 or 6 for each particle amount and poison percentage (**Supplementary Fig. S1**; **Table 1**).

#### Hb

A total of 360 3-D classification jobs using a combination of 2, 3, or 5 classes with varying T values of 1, 2, 4, and 8 for each particle amount and poison percentage (**Supplementary Fig. S2**; **Table 1**).

#### mortalin-GrpEL1

A total of 216 3-D classification jobs using a combination of 2, 3, or 4 classes with varying T values of 1, 3, and 7 for each particle amount and poison percentage (**Supplementary Fig. S2**; **Table 1**).

#### MoFeP

A total of 144 3-D classification jobs using a combination of 2, 3, or 5 classes with varying T values of 1, 2, 4, and 8 for each particle amount and poison percentage (**Supplementary Fig. S3**; **Table 1**).

#### T20S

A total of 108 3-D classification jobs using a combination of 2, 3, or 4 classes with varying T values of 1, 4, and 20 for each particle amount and poison percentage (**Supplementary Fig. S3**; **Table 1**).

### Feature engineering from 3-D classification metadata within RELION

To classify particles based on their behavior in 3-D classification, we extract statistical metadata from RELION’s 3-D particle data STAR files to use as input features. Specifically, we utilize per-particle metadata generated over the course of 3-D classification jobs comprising 25 expectation-maximization (EM) iterations. All iterations, excluding the zeroth and first (which typically contain noisy data before the classification begins to converge), are parsed and processed using the ANTIDOTE framework. These resulting features capture particle class assignments, likelihood statistics, and classification stability information over multiple iterations. Once the raw classification data are parsed, we apply a series of feature engineering techniques to aggregate, encode, and normalize these statistics into a format suitable for training the ANTIDOTE neural network (**Fig. 3a**).

During 3-D Classification in RELION, each particle in a 3-D Classification job is assigned to a class at each iteration based on probability distributions computed during EM refinement.^8,13^ We extract the following statistical features reported by RELION that describe how each particle behaves over classification iterations:

#### Log-Likelihood Contribution (rlnLogLikeliContribution)

Measures the per-particle contribution to the likelihood of its assigned class. Higher values indicate a well-classified particle with a large contribution whereas lower values suggest uncertainty or noise.

#### Maximum Value in the Probability Density (rlnMaxValueProbDistribution)

Denotes the highest posterior probability value across all class distributions for a given particle, reflecting the strength of classification certainty. Higher values indicate greater classification certainty, whereas lower values suggest ambiguity in class assignment.

#### Number of Significant Samples (rlnNrOfSignificant-Samples)

Quantifies the number of non-trivial peaks in the particle’s class probability distribution. Higher counts may reflect classification ambiguity, especially in the presence of heterogeneity or low-quality input data.

Across all extracted RELION 3-D metadata, we apply five feature transformations to increase the dimension of our input vector to better represent particle behavior:

#### Iteration Class Frequency Encoding

For each particle *i* at iteration *k*, we compute the fraction of total particles assigned to the same class *C*_*i,k*_. This captures both temporal and global class distribution dynamics. With *P* denoting the total number of particles and *I*(°) denoting the identity function, equal to 1 if the argument is true and 0 otherwise, the encoded frequency feature *F* (*i, k*) is defined as (Equation 1):

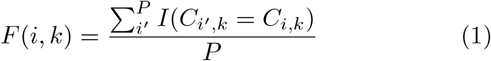

#### Sum of Class Transitions Over Iterations

To capture classification instability over time, we compute the number of class transitions per particle across at each iteration up to that iteration, where N=25 is the total number of iterations (Equation 2):

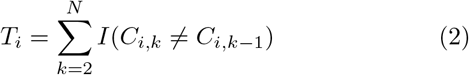

#### Target Encoding of Class Assignments to Fourier Completeness and Resolution

Instead of treating class assignments as categorical numbers, we map each class to two physically meaningful properties derived from RELION model metadata: Fourier completeness (rlnOverallFourierCompleteness), representing how well the assigned class covers frequency space in the final 3-D reconstruction, and estimated resolution (rlnEstimatedResolution), indicating the estimated spatial resolution achieved for a given class. These values are then assigned to each particle based on its class at each iteration.

#### Z-score Normalization of Numerical Features

For each iteration, we normalize the distributions of rlnLogLikeliContribution, rlnMaxValueProbDistribution, and rlnNrOfSignificantSamples across all particles using Z-score normalization. These normalized values are included as additional features to control for scale variability and promote stable learning.

Following the application of these transformations, we aggregate the engineered features into a final dataset. The resulting feature vector for each particle comprises 241 dimensions: 24 values for each of 10 iterative features (covering 24 iterations), plus one global value for the unique class change count, computed across all iterations (10 × 24 + 1 = 241). Each feature is subsequently mean-normalized across particles and iterations to facilitate efficient model convergence during training. The full dataset is stored in HDF5 format, which enables efficient, chunk-wise streaming during model training and tuning, avoiding the need to load the entire dataset into memory. All code for parsing, engineering, and normalizing RELION 3-D Classification metadata is available on GitHub and accessible via the ‘antidote train’ command. HDF5 files used for training are available from the authors upon request.

### Neural network model architecture, training, and evaluation

ANTIDOTE is a fully-connected multilayer perceptron (MLP) that predicts the likelihood of a particle being deleterious. Formally, the model defines a function (Equation 3):

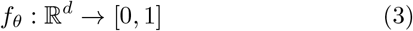

where *x* ∈ ℝ^*d*^ is a feature vector representing a particle, and *ŷ* = *f*_*θ*_(*x*) is the predicted probability that the particle is deleterious. The model is optionally preceded by a Multi-Head Self-Attention (MHA) encoder that allows the network to learn pairwise feature interactions.^30^ All layers use Keras default weight initializers; dense weights are initialized via Glorot uniform and biases with zeros.^49^ Training is performed using the Adam optimizer to minimize binary cross-entropy loss.^50^

Performance was evaluated on held-out validation and test sets using a comprehensive suite of metrics: AUROC, AUPRC, binary cross-entropy, Brier score, accuracy, precision, recall, balanced accuracy, and false discovery rate. Metrics were computed after each chunked training step and persisted to JSON logs for reproducibility and fine-grained analysis. Generally, training and validation were performed on 5 out of the 6 proteins in the training data set described above using an 80:20 split; and testing was performed on the 6^*th*^ protein. During model architecture evaluation and dataset analysis, models were often trained and validated on 2 proteins and tested on 4 (**Fig. 3a, Table 1**).

For hyperparameter optimization, we employ Keras Tuner Hyperband with the objective of maximizing validation AUROC.^31^ The search space includes the use of batch normalization, the L2-regularization strength (*λ* ∈ [10^−5^, 10^−2^]), optimizer learning rate ([10^−4^, 10^−2^] log-uniform), dropout rate (0.0–0.4), and batch size (256, 512). When MHA is enabled, the number of attention heads (2–8) and key dimension size (16–64) are also included in the tuner search space. To generate a final model instance from a training run, the top-ranked model from the Hyperband search is reinitialized and retrained for 20 epochs with early stopping, using chunked streaming from HDF5 to avoid loading the full dataset into memory.

The most performant model employs a 4-head MHA layer with a key dimension of 32, followed by three fully-connected layers with 241, 241, and 120 units respectively. All dense layers use ReLU activations, batch normalization, and L2 regularization with strength *λ* = 0.00197; without dropout. The model was trained with a learning rate of 0.00071, using a batch size of 256 and a final sigmoid output unit for binary classification. We note that many of the models generated by our training runs were comparable in terms of accuracy for each protein. The exclusion of Multi-Head Attention results in models with only a marginal decrease in performance with faster inference time. Performant examples of both model architectures are included with ANTIDOTE as ‘antidote-main’ and ‘antidote-mini’ respectively.

ANTIDOTE is implemented in TensorFlow.^51,52^ Data preparation and visualization are performed with tools from the SciPy software stack, as well as invaluable tools from the teamtomo ecosystem.^53–58^

### Benchmarking of ANTIDOTE using EMPIAR datasets

#### EMPIAR-12743

Raw micrographs (2368 movies; 4096 x 4096 pixels, 0.735 Å/pixel, 60 electrons/Å^2^) were imported into cryoSPARC for patch-based motion correction and patch CTF estimation. Particles were selected using a template-based particle picker using EMD-458159 as a reference volume. Selected particles were then extracted with a 256-pixel box and Fourier cropped to a box size of 64 pixels (2.94 Å/pixel). These initial picks (2,444,215 particles) were subjected to *ab initio* (1 class, C_1_ symmetry) reconstruction to generate a volume that was then subjected to a non-uniform refinement (C_2_ symmetry). These particles are hereafter referred to as “uncurated” particles. To generate a “standard” processing workflow, all 2,444,215 initial particle picks were re-extracted at a 264-pixel box size (0.735 Å/pixel) and subjected to a non-uniform refinement (C_2_ symmetry) with CTF refinement using the ab-initio volume as a reference to yield a 4.06 Å resolution reconstruction. To generate an “ANTIDOTE-curated” workflow, we exported the initial binned particle stack (2,444,215 particles; 64 pixels (2.94 Å/pixel)) to RELION-5.0 using the csparc2star from pyem and performed a 3-D classification (K=5, T=1). The outputs were then subjected to inference by ANTIDOTE and particles identified as constructive by ANTIDOTE were imported back into cryoSPARC. These particles were then re-extracted with a 256-pixel box size (0.735 Å/pixel) and subjected to a final non-uniform refinement (C_2_ symmetry) with CTF parameters and aberration refinements to yield a 2.70 Å resolution reconstruction (**Figure 5a**).

To generate a “partially curated” particle stack, we performed one round of 2-D classification on the uncurated particle stack and discarded any obvious non-protein classes. We exported the resulting particle stack (788,417 particles) to RELION-5.0 using csparc2star from pyem and performed the same 3-D classification (K=5, T=1) followed by inference with ANTIDOTE as described above. The ANTIDOTE-curated particle stack (693,178 particles with an ANTIDOTE assignment of 0.5 or higher) was then imported into cryoSPARC and a non-uniform refinement (C_2_ symmetry) was performed. The particles were then extracted at a 256-pixel box size (0.735 Å/pixel) and a final non-uniform refinement (C_2_ symmetry) with CTF and aberration refinements to yield a 2.60 Å resolution reconstruction. For comparison, the selected 2-D class particles were subjected to the same series of non-uniform refinements, yielding a 2.79 Å resolution reconstruction (Figure 5b).

The final “curated” particle stack was generated by abinitio model generation (3 class, C_1_ symmetry) and the particles contributing to the two classes most resembling Hb were selected. These particles were subjected to a non-uniform refinement (C_2_ symmetry) and then exported to RELION-5.0 using csparc2star from pyem where the same 3-D Classification (K=5, T=1) and inference by ANTIDOTE were performed as described above. The resulting ANTIDOTE-curated particles (219,746 particles) were then imported into cryoSPARC. These particles were then re-extracted with a 256-pixel box size (0.735 Å/pixel) and a final non-uniform refinement (C_2_ symmetry) with CTF and aberration refinements was performed to yield a 2.71 Å resolution reconstruction. For comparison, the selected ab-initio particles (225,123) were subjected to the same non-uniform refinement (C_2_ symmetry) and extraction protocol as the ANTIDOTE-curated particles to yield a 2.73 Å resolution reconstruction (**Figure 5c**).

#### EMPIAR-48384

Raw micrographs (2000 movies; 4096 x 4096 pixels, 0.735 Å/pixel, 60 electrons/Å^2^) were imported into cryoSPARC for patch-based motion correction and patch CTF estimation. Particles were selected using a blob picker (100 Å inner and 200 Å outer particle diameter). These particles were extracted with a 96-pixel box size (2.94 Å/pixel). These particles were then subjected to a single round of 2-D classification where only classes with obvious protein features were selected. An initial volume was generated using ab initio model generation (1 class, C_1_ symmetry) in cryoSPARC using the selected particles. These particles were then exported to RELION-5.0 and a 3-D classification (K=5, T=4) was performed. Particles from the resulting 3-D classification were then subjected to inference by ANTIDOTE. The selected ANTIDOTE particles were then imported back into cryoSPARC and extracted with a 384-pixel box (0.735 Å/pixel) and subjected to a final non-uniform refinement (C_2_ symmetry) with CTF and aberrations refinements to yield a 2.41 Å resolution reconstruction. To provide a comparison to ANTIDOTE curation, the 2-D-selected particles were extracted at a 384-pixel box (0.735 Å/pixel) and subjected to a non-uniform refinement (C_2_ symmetry) with CTF and aberration refinements to yield a 2.49 Å resolution reconstruction (**Supplementary Fig. S10**).

#### EMPIAR-11304

Raw micrographs (783 movies; 5760 x 4092 pixels, 0.87 Å/pixel, 40 electrons/Å^2^) were imported into cryoSPARC for patch-based motion correction and patch CTF estimation. Particles were identified using a blob picker (140 Å inner and 160 Å outer particle diameter)and these particles were then extracted with a 128-pixel box size (1.74 Å/pixel). The initial model for the 3-D classification was generated by using the starting particle stack in cryoSPARC and running ab initio model generation (3 classes, D_7_ symmetry) with the best class being used for 3-D Classification. The blob-picked particles were then exported to RELION-5.0 for 3-D classification (K=5, T=4). Particles from the resulting 3-D classification was then either subjected to inference by ANTIDOTE or those contributing to the best classes (e.g., classes 1 and 3), chosen based on volume completeness and quality, were combined. Both the ANTIDOTE-selected particles as well as the particles in classes 1 and 3 were imported back into cryoSPARC for reference-based motion correction. These selected particle stacks were each subjected to a non-uniform refinement with CTF and aberration refinements, yielding a 2.58 Å resolution reconstruction for the ANTIDOTE stack and 2.89 Å resolution reconstruction for the class 1/3 combined stack (**Supplementary Fig. S11**).

#### EMPIAR-11668

Raw micrographs (3,285 movies; 5760 x 4092 pixels, 0.86 Å/pixel, 60 electrons/Å^2^) were imported into cryoSPARC for patch-based motion correction, patch CTF estimation, and denoising. Initial particle coordinates were then identified from the denoised micrographs using a blob picker and were extracted at a 64-pixel box size (3.44 Å/pixel). Ab-initio (1 class, C_1_ symmetry) was then performed to generate a starting model. These particles were then exported to RELION-5.0 using csparc2star and a 3-D Classification (K=5, T=4) was performed (10 Å low-pass filter, D_2_ symmetry). To generate a starting volume for 3-D classification, the initial particle stack in cryoSPARC was subjected to 3 rounds of 2-D classification where obvious protein density was selected. Inference was then run on the resulting 3-D classification job and particles above a cutoff of 0.5 were selected and re-imported into cryoSPARC. Additionally, class 5 of the 3-D classification was selected as a control an imported into cryoSPARC. Both particle stacks were then run through separate ab initio model generation jobs (single class, C_8_ symmetry applied) and the two best volumes resembling proteinaceous density were selected. A final non-uniform refinement was performed followed by local resolution estimation (**Supplementary Fig. S12**).

## Author contributions

M.A.H. conceived the project, defined its overall scope, and supervised all aspects of the study. R.F.B., D.J., and A.F. developed the core ANTIDOTE codebase and implemented training and testing approaches for ANTIDOTE models. B.D.C. generated the ground-truth labeled particle datasets, with support from M.J.B. and Y.H., performed extensive 2-D and 3-D classifications, and conducted associated analyses. R.F.B., B.D.C., D.J., and M.A.H. analyzed and interpreted the data, prepared figures, and wrote the manuscript. All authors discussed the results and contributed to manuscript revision.

## Acknowledgements

We are grateful to Prof. Kevin Corbett for providing valuable feedback on this manuscript, and the entirety of the Herzik lab for facilitating insightful discussions. The authors acknowledge Brendan Dennis, Kevin Smith, and the UCSD Physics Computing Facility for their insights and support. Molecular graphics and analyses were performed with UCSF ChimeraX, developed by the Resource for Bio-computing, Visualization, and Informatics at the University of California, San Francisco (UCSF), with support from National Institutes of Health (NIH) grant R01-GM129325 and the Office of Cyber Infrastructure and Computational Biology, National Institute of Allergy and Infectious Diseases. This work was funded by the NIH grants R35-GM138206 (M.A.H), T32-GM008326 (M.J.B), and S10-OD032471 (M.A.H). Additional support includes the Searle Scholars Program (M.A.H.), a George W. and Carol A. Lattimer UCSD Faculty Research Fellowship (M.A.H), and a Cottrell Scholars Award (M.A.H.). B.D.C is partly supported by Goeddel Family Technology Sandbox Fellowship.

## Competing interests

The authors declare that they have no competing interests.

## Data availability

ANTIDOTE is available at https://github.com/herzik-lab/ANTIDOTE/

## Notes

### Competing Interest Statement

The authors have declared no competing interest.

### Summary of Updates

The revised manuscript fixes a formatting issue.

