## Supplemental Files for "ANTIDOTE: A Metadata-Driven Neural Network for Improving CryoEM 3-D Particle Sorting"

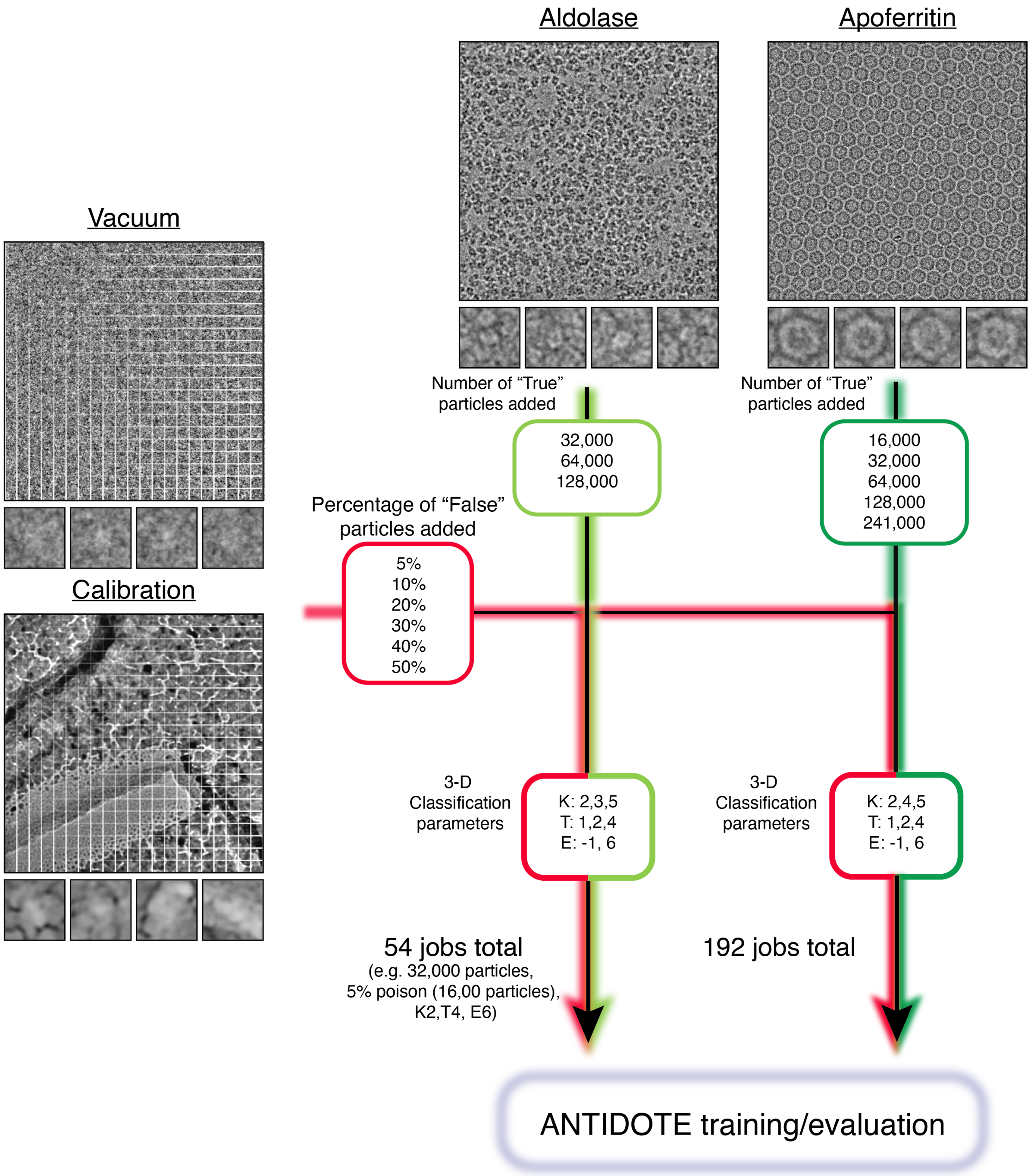


**Figure S1: Generation of poisoned aldolase and apoferritin datasets.** (*Top*) Aligned micrographs of aldolase (light green) and apoferritin (dark green) are presented with representative particle picks shown below. (*Left*) Aligned micrographs of images taken over vacuum (pink) or of a crossed-lines replica grating calibration grid (red) are presented with grid-like "particle" spacing shown as white lines and representative "particle" picks shown below. The particle counts from each dataset used for combination as well as the total number of 3-D classification jobs performed are annotated.

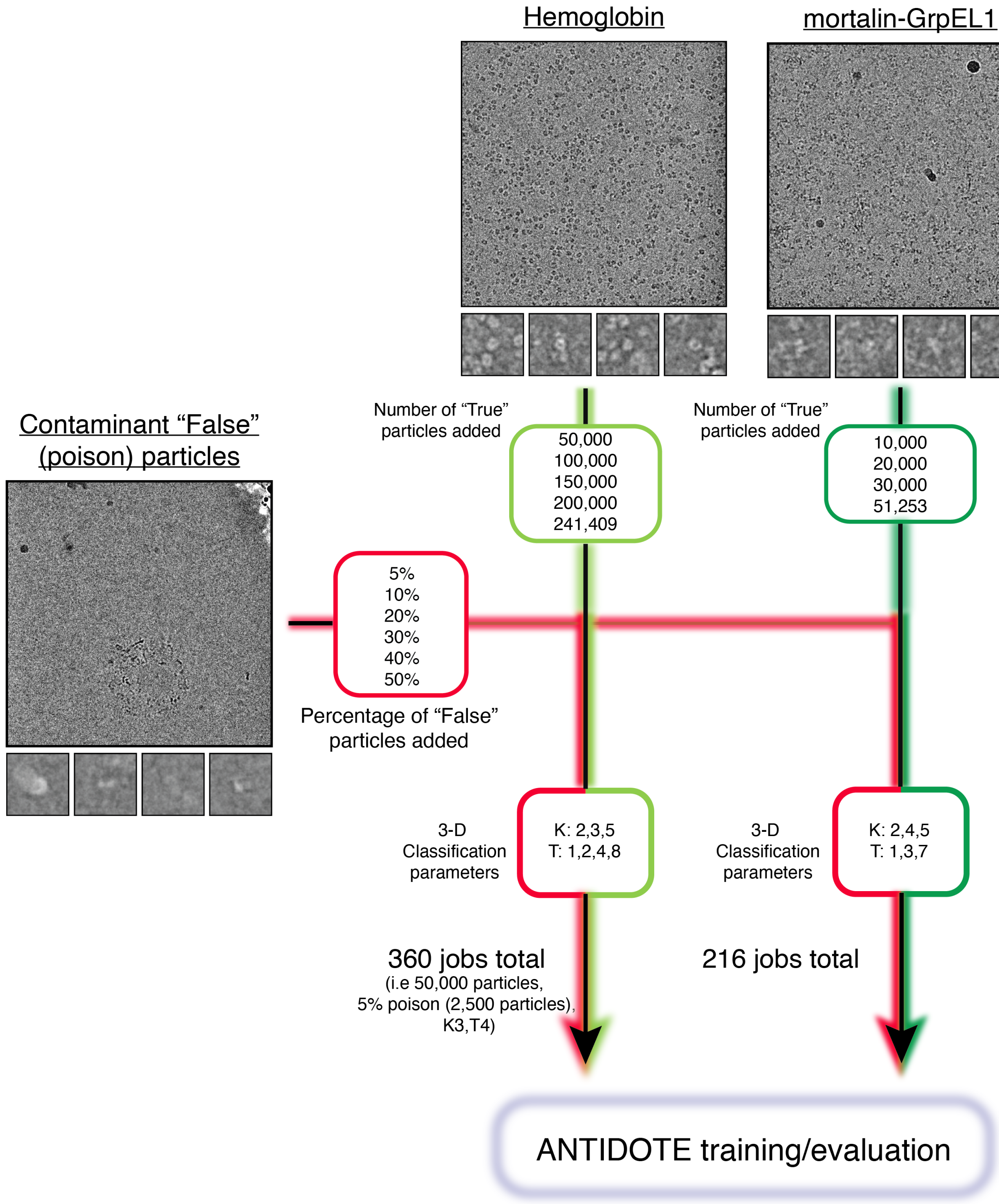


**Figure S2: Generation of poisoned hemoglobin and mortalin-GrpEL1 datasets.** (*Top*) Aligned micrographs of hemoglobin (light green) and mortalin-GrpEL1 (dark green) are presented with representative particle picks shown below. (*Left*) Aligned micrographs of images taken of contaminant particles (red) are presented with representative “particle” picks shown below. The particle counts from each dataset used for combination as well as the total number of 3-D classification jobs performed are annotated.

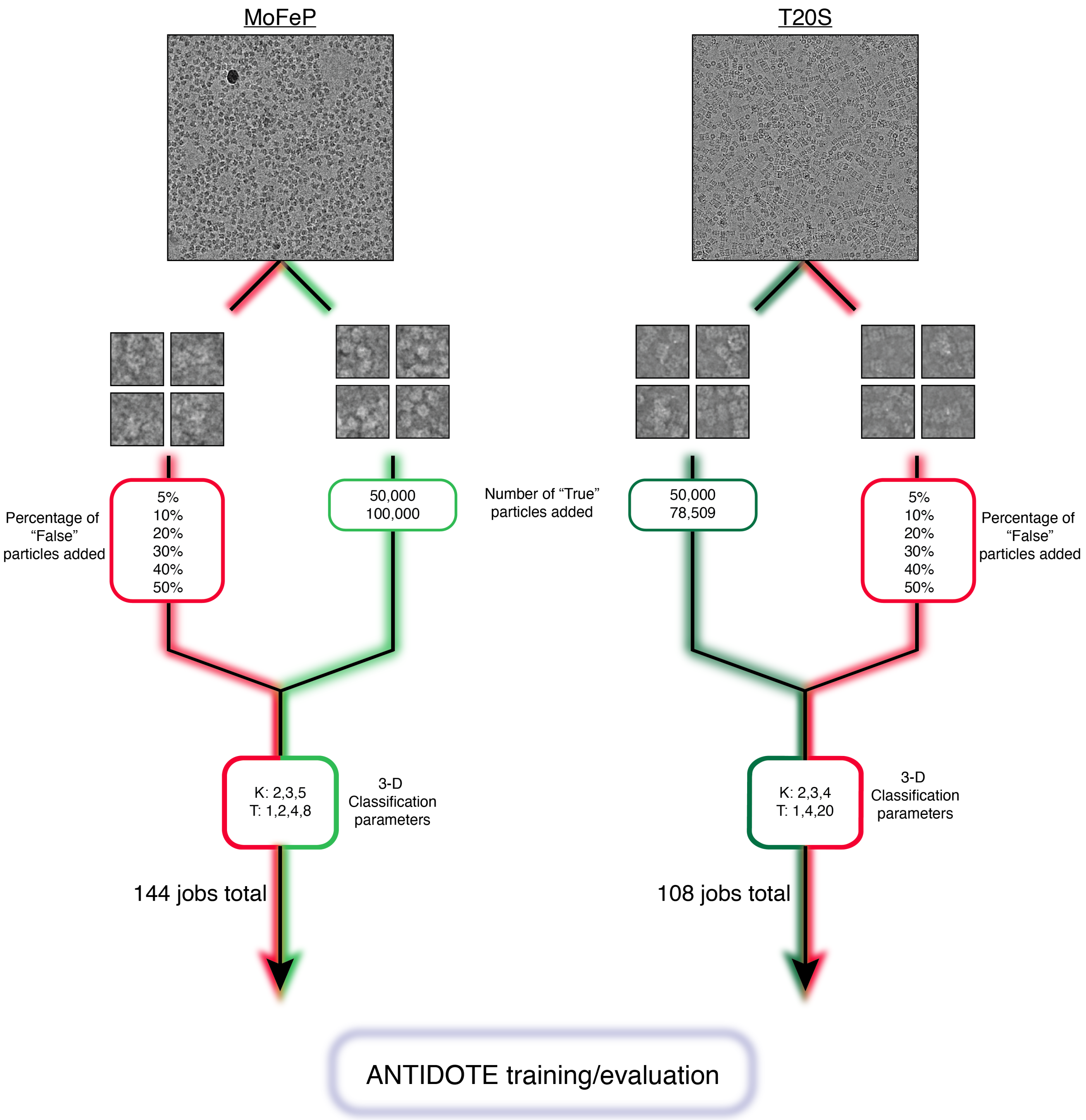


**Figure S3: Generation of poisoned MoFeP and T20S datasets.** (*Left*) Aligned micrograph of MoFeP is presented with "True" particle (light green) and poison particle (red) picks shown below. (*Right*) Aligned micrograph of T20S proteasome core is presented with "True" particle (green) and "False" (poison) particle (red) picks shown below. The particle counts from each dataset used for combination as well as the total number of 3-D classification jobs performed are annotated.

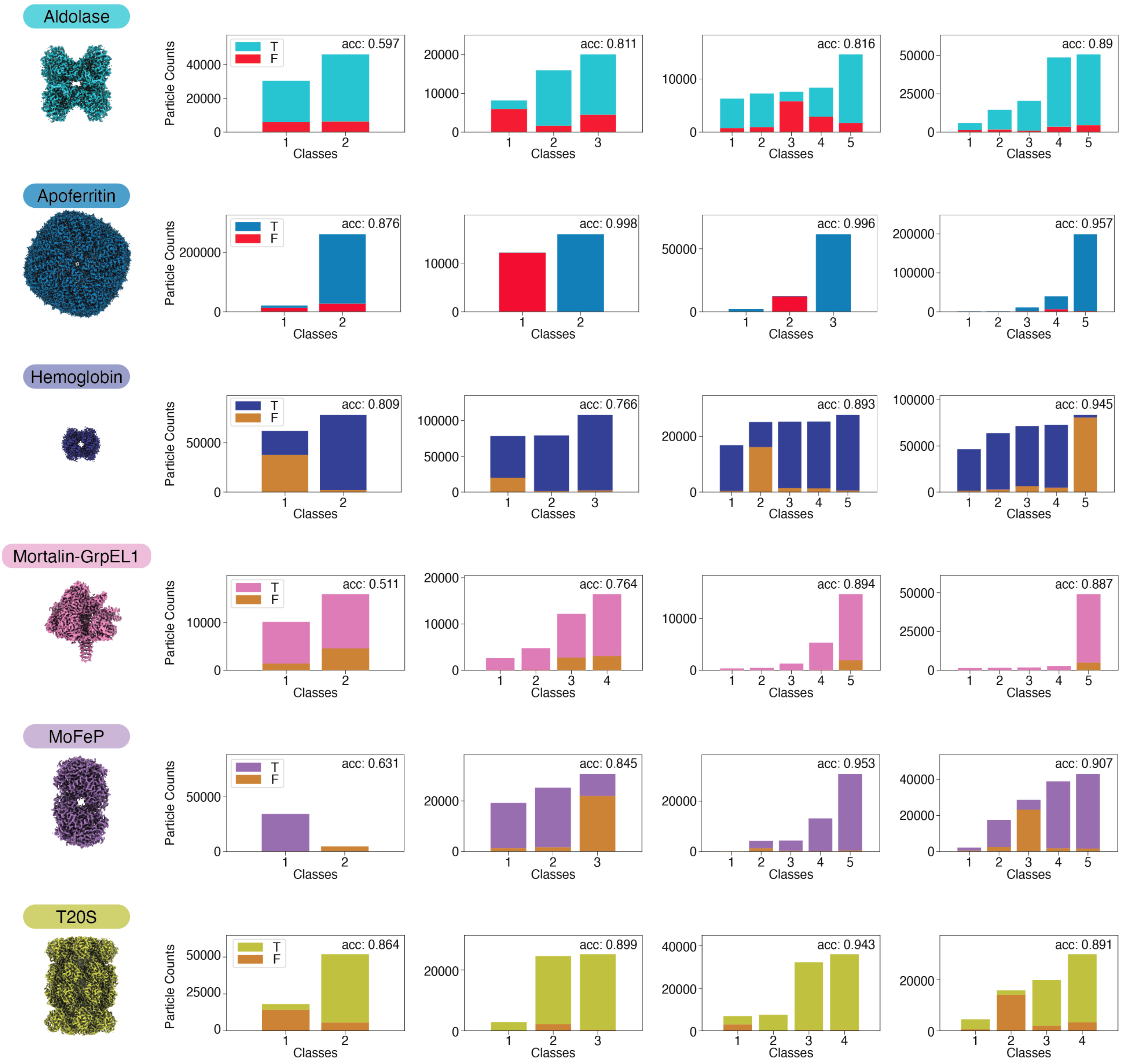


**Figure S4: Example 3-D Classification results with ground-truth labels.** Stacked bar plots representing the class composition for 3-D classification runs for each of the six proteins in our panel. Bar plots are colored by ground-truth label. Accuracies are calculated for the best combination of accepted and rejected classes while enforcing the rejection of at least one class, as described in the Results.

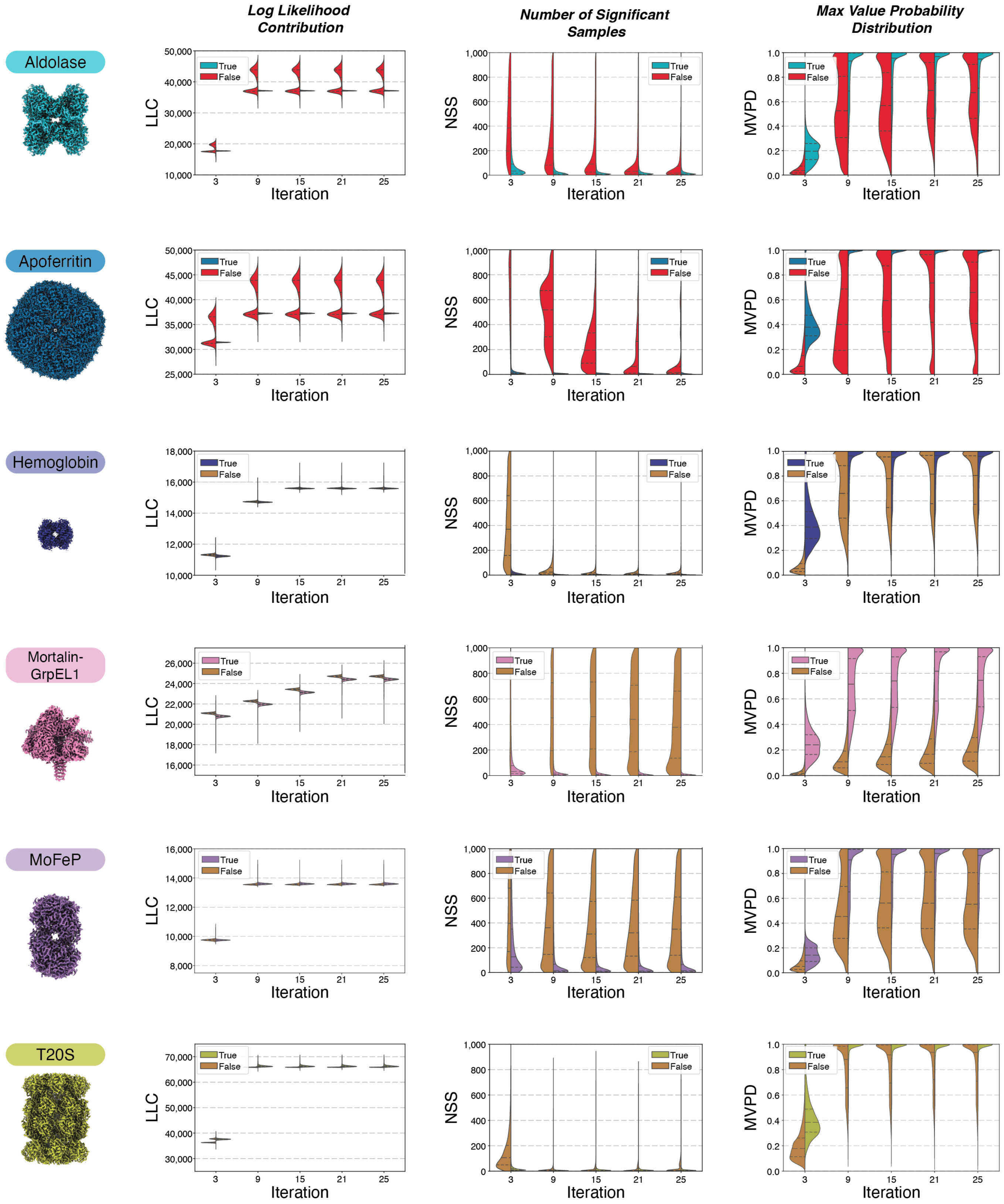


**Figure S5: 3-D Classification metadata analyses of each protein sample set.** For each protein sample assessed in this study, a 3-D reconstruction of the protein sample (*left*) and violin plots of log-likelihood contribution (LLC), number of significant samples (NSS), and maximum value in the probability distribution (MVPD) across iterations for True and False particles are shown.

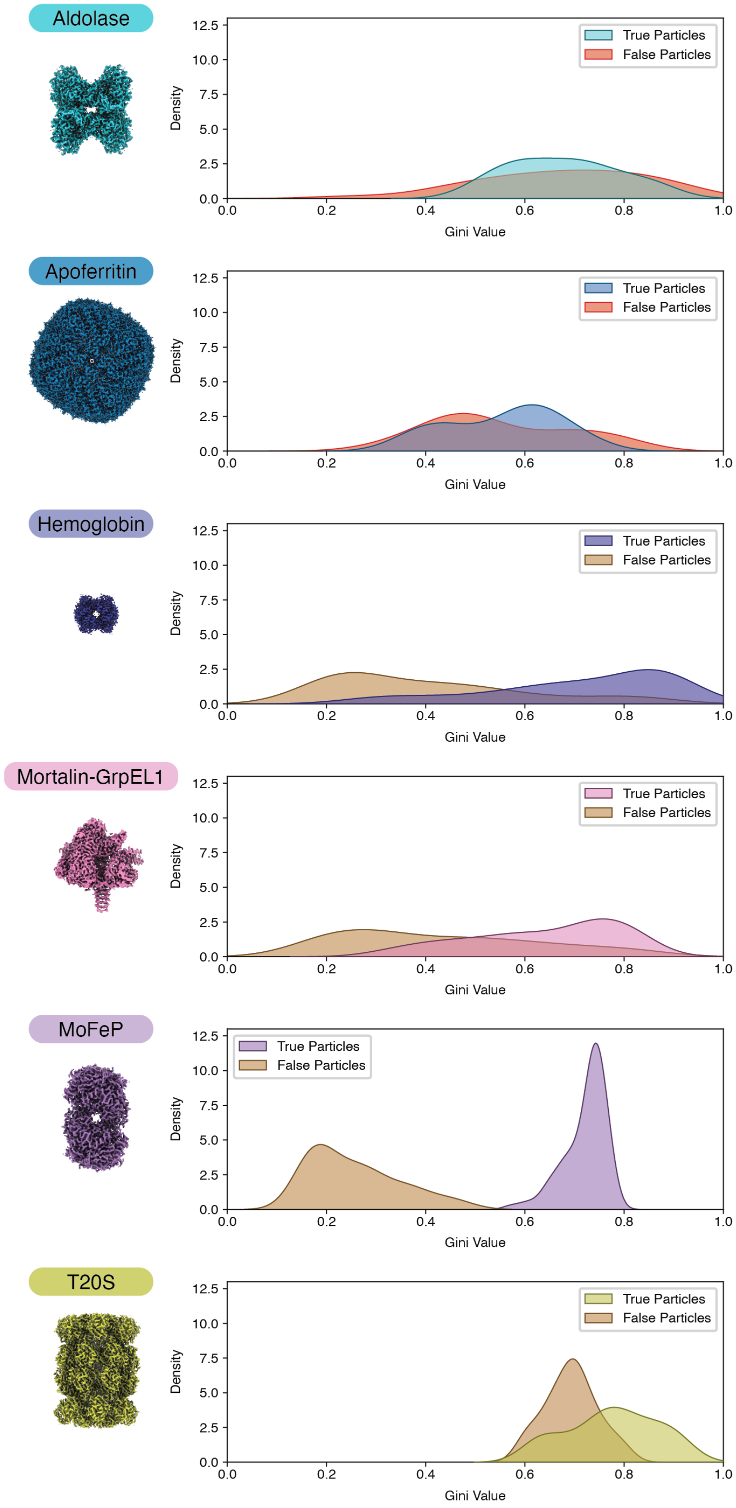


**Figure S6: Gini value distributions for each protein sample set from 3-D Classification.** For each protein sample assessed in this study, a 3-D reconstruction of the protein (*left*) is shown with Gini distributions for True- and False-labeled particles across all 3-D classifications assessed using the ground-truth-labeled evaluation datasets.

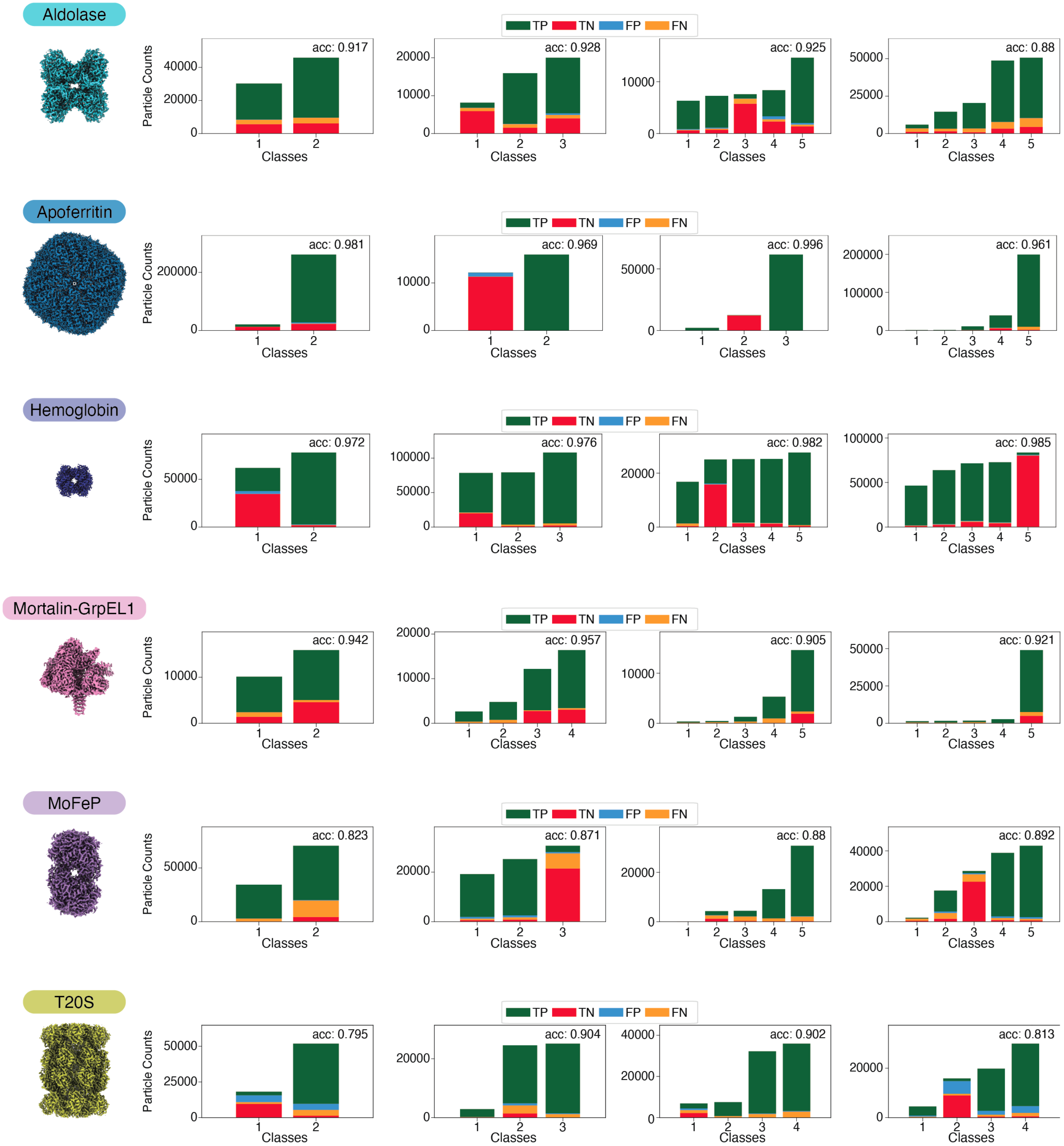


**Figure S7: Example ANTIDOTE predictions for 3-D Classifications run with ground-truth labels.** Stacked bar plots representing the class composition for 3-D classification runs for each of the six proteins in our panel, labeled with predictions generated by ANTIDOTE inference. Bar plots are colored by the confusion matrix values generated using the ANTIDOTE-derived labels, with ground-truth labels acting as an oracle. Accuracies are calculated for the ANTIDOTE predictions. All 3-D classifications are identical to those represented in **Figure S4**.

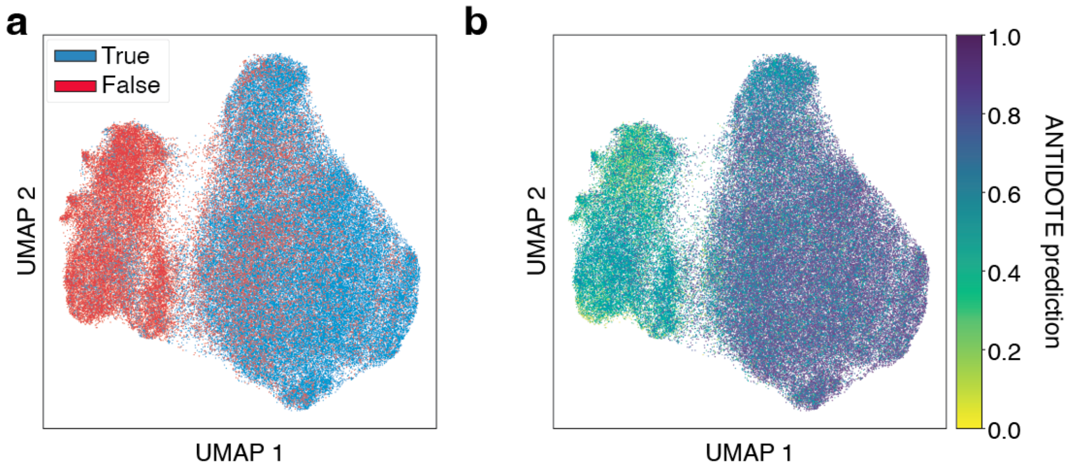


**Figure S8: Comparison of ANTIDOTE prediction versus cryoDRGN classification using T20S proteasome core ground-truth labels. a)** CryoDRGN latent space UMAP projection colored by CryoSPARC-derived labels for poisoned T20S proteasome. **b)** Same CryoDRGN latent space UMAP projection as (**a** colored by ANTIDOTE-derived predictions.

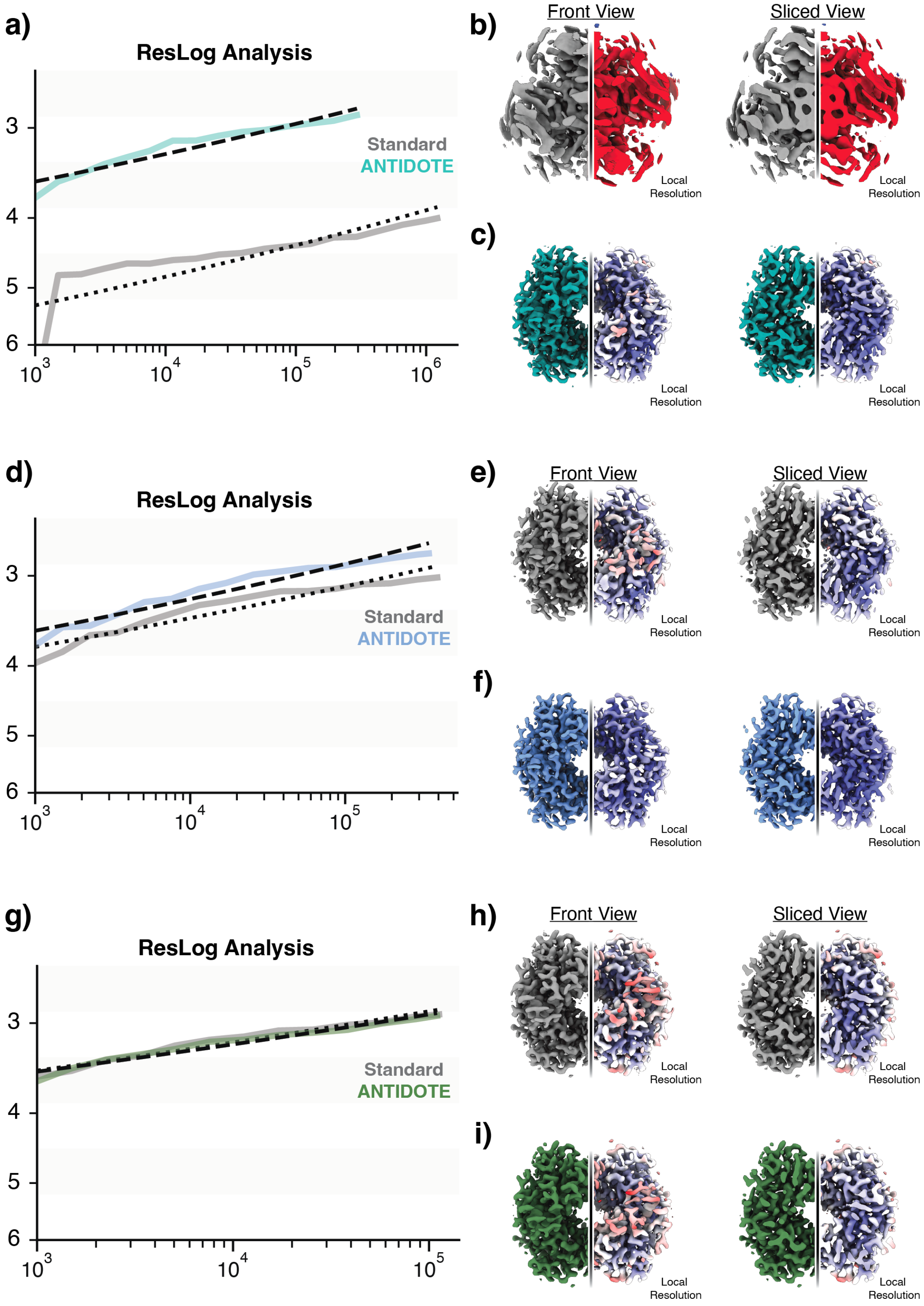


**Figure S9: ANTIDOTE improves particle efficiency and enhances global and local resolution at early stages of particle curation. a,d,g)** Resolution versus particle number (ResLog) analyses comparing standard particle curation workflows (gray/black) to ANTIDOTE-based workflows (teal, light blue, or blue) at three stages of traditional particle curation. ANTIDOTE curation consistently achieves higher resolution with fewer particles and displays steeper ResLog slopes, indicating improved particle quality and reconstruction efficiency. **b,c,e,f,h,i)** Corresponding local resolution maps of final 3-D reconstructions obtained from traditional (**b,e,h**) and ANTIDOTE-curated (**c,f,i**) datasets for each target. For each refinement, front and sliced views through the 3-D reconstructions (*left*) and volumes colored by local resolution estimates (*right*) reveal globally and locally improved map quality.

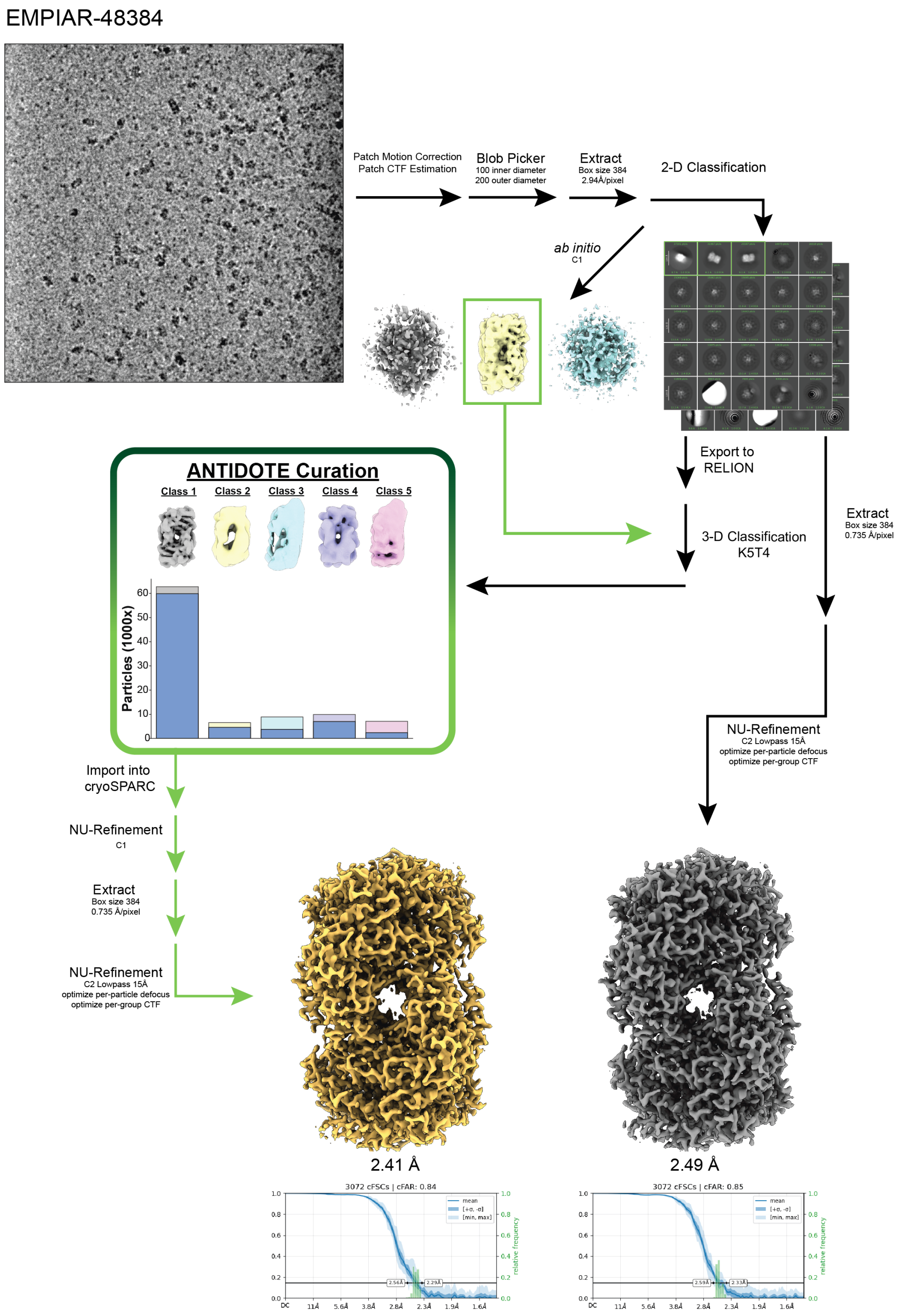


**Figure S10: Single-particle cryoEM data processing and ANTIDOTE curation workflow for EMPIAR-48384 (MoFeP).** Representative aligned and denoised micrograph from EMPIAR-48384 (MoFeP) showing particle density and distribution. Following motion correction, CTF estimation, and micrograph denoising, particles were identified using blob picking, extracted, and subjected to successive rounds of 2-D classification in cryoSPARC. Particle stacks were exported to RELION for 3-D classification into five major classes (Classes 1–5). True and False particle distributions are shown below each class. Particles from each class were curated using the ANTIDOTE metadata-based classifier. Curated particle subsets were re-imported into cryoSPARC for *ab initio* reconstruction and subsequent non-uniform refinement, including per-particle CTF refinement and optimization of defocus parameters. Final reconstructions refined to 2.41 Å (ANTIDOTE-curated) and 2.49 Å (Traditionally-curated) resolution. Fourier shell correlation (FSC) curves and 3-D FSC plots following non-uniform refinement are shown for each map, highlighting the improved directional resolution and isotropy obtained after ANTIDOTE-based particle selection.

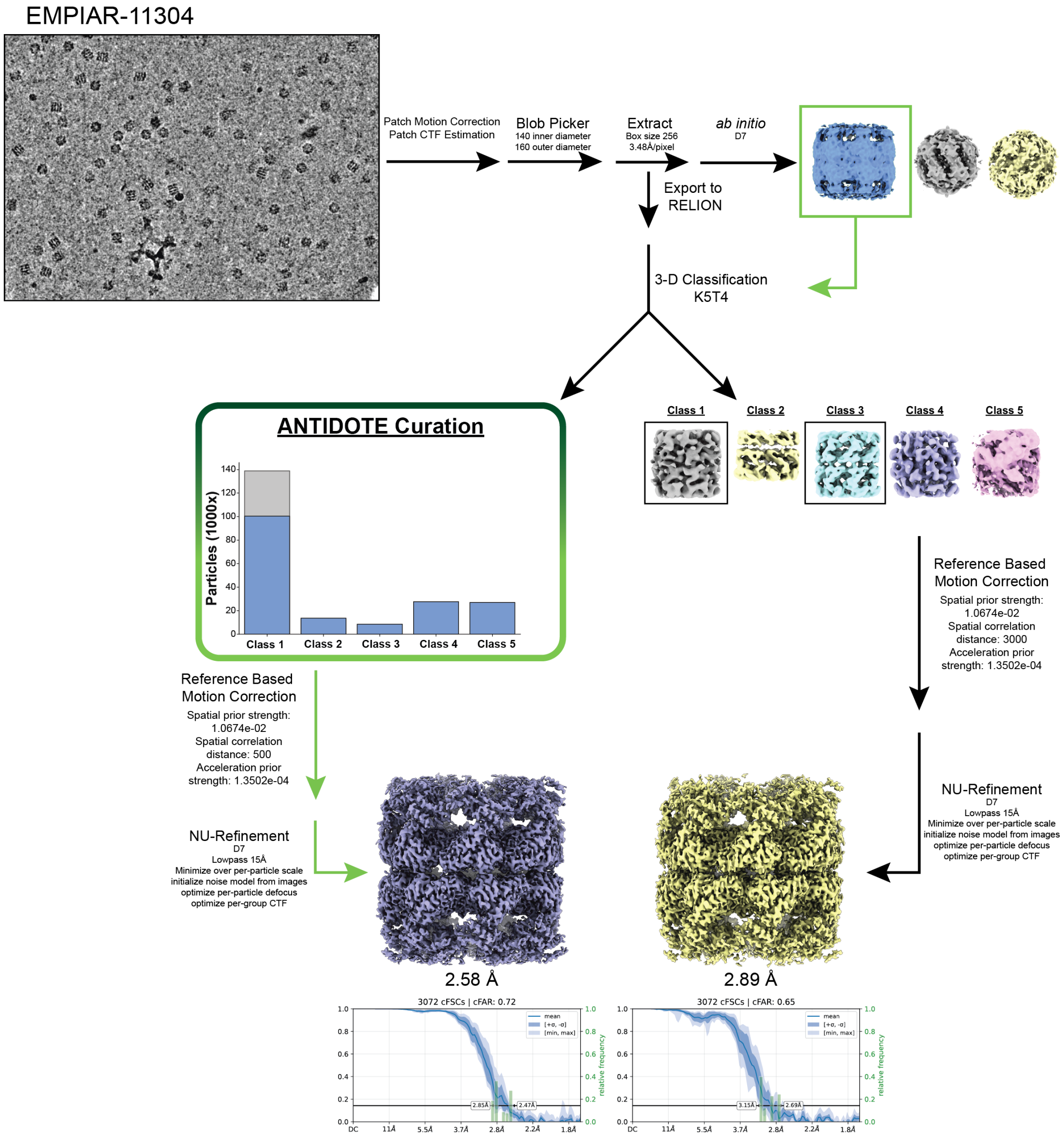


**Figure S11: Single-particle cryoEM data processing and ANTIDOTE curation workflow for EMPIAR-11304 (GroEL).** Representative aligned and denoised micrograph from EMPIAR-11304 (GroEL) illustrating particle density and distribution. Following motion correction, CTF estimation, and micrograph denoising, particles were identified using blob picking, extracted, and subjected to successive rounds of 2-D classification in cryoSPARC. Particle stacks were exported to RELION for 3-D classification into five major classes (Classes 1–5). True and False particle distributions are shown below each class. Particles from each class were curated using the ANTIDOTE metadata-based classifier. Curated particle subsets were re-imported into cryoSPARC for *ab initio* reconstruction and subsequent non-uniform refinement, incorporating reference-based motion correction to improve per-particle alignment. This yielded final reconstructions at 2.58 Å (ANTIDOTE-curated) and 2.89 Å (Traditionally-curated) resolution. Fourier shell correlation (FSC) curves and 3-D FSC plots following non-uniform refinement are shown for each final map.

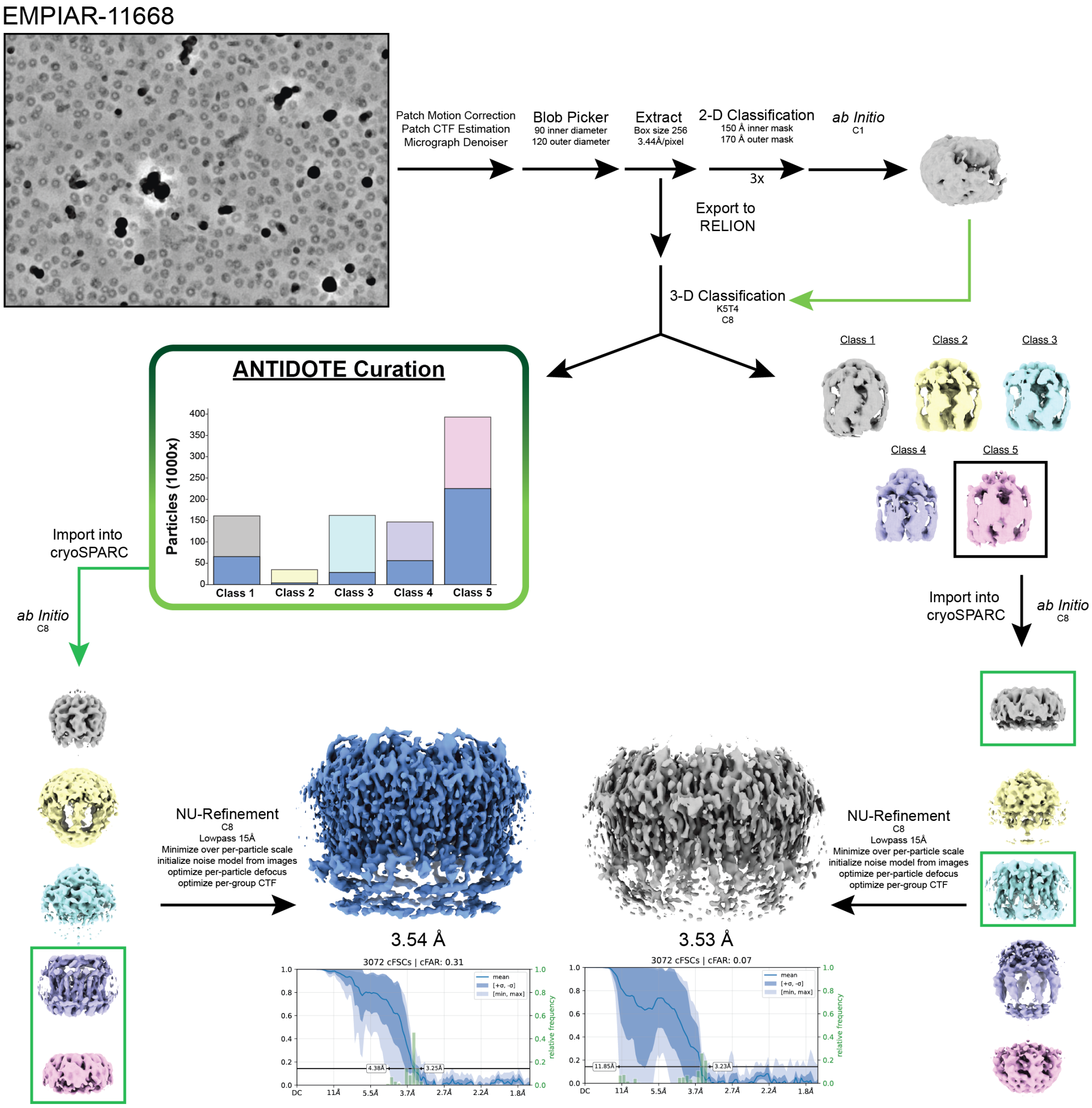


**Figure S12: Single-particle cryoEM data processing and ANTIDOTE curation workflow for EMPIAR-11668 (CALHM1).** Representative aligned and denoised micrograph from EMPIAR-11668 demonstrating particle density and distribution. Following motion correction, CTF estimation, and micrograph denoising, particles were identified using blob picking, extracted, and subjected to successive rounds of 2-D classification in cryoSPARC. Resulting particle stacks were exported to RELION for 3-D classification into five major classes (Classes 1–5). True and False particle distributions are shown below each class. Particles from each class were curated using the ANTIDOTE metadata-based classifier. Curated particle subsets were re-imported into cryoSPARC for *ab initio* reconstruction and non-uniform refinement, producing final reconstructions at 3.54 Å (ANTIDOTE-curated) and 3.53 Å (Traditionally-curated) resolution. Fourier shell correlation (FSC) curves and 3-D FSC plots following non-uniform refinements are shown for each refined map.
